# The population structure and genetic health of European wolves

**DOI:** 10.64898/2026.03.20.712003

**Authors:** Evelyn T. Todd, Claudia Fontsere, Xin Sun, Camilla Hjorth Scharff-Olsen, Germán Hernández-Alonso, Liam Thomas Lanigan, Nuno Filipe Gomes Martins, Marta Maria Ciucani, Jazmín Ramos-Madrigal, Lauren Hennelly, Sarah S.T. Mak, Zanete Andersone-Lilley, Angelica Åsberg, Linas Balčiauskas, Laima Baltrūnaitė, Gennady F. Baryshnikov, Bazartseren Boldbaatar, Bazartseren Boldgiv, Barbora Černá Bolfíková, Tomasz Borowik, Dominika Bujnáková, Paolo Ciucci, Ayşegül Karaahmetoğlu Çoban, Emrah Çoban, Maria Erlandsson, Øystein Flagstad, Laurent Frantz, Eli Geffen, Jenni Harmoinen, Lina Jelk, Daniela C. Kalthoff, Alexandros Α. Karamanlidis, M. Çisel Kemahlı-Aytekin, Ilpo Kojola, Alexander Kopatz, Pavel Kosintsev, Josip Kusak, Anastasiia Kuznetsova, Laura Kvist, Linn Larsson, Greger Larson, Anna Linderholm, Frode Lingaas, Luis Llaneza, Sarah L. F. Martin, Shai Meiri, Øyvind Øverli, Ladislav Paule, Mikhail Sablin, Morten Skage, Carola Stålfjäll, David W.G. Stanton, Çağan H. Şekercioğlu, Renata Špinkyte, Konstantin Tirronen, Linda Törngren, Cristiano Vernesi, Nobuyuki Yamaguchi, Mikael Åkesson, Robert W. Mysłajek, Pavel Hulva, Carsten Nowak, Magdalena Niedziałkowska, Sabina Nowak, Love Dalén, Raquel Godinho, Urmas Saarma, Luca Fumagalli, Jouni Aspi, Hans K. Stenøien, Michael D. Martin, Mikkel-Holger S. Sinding, M. Thomas P. Gilbert, Shyam Gopalakrishnan

## Abstract

Once nearly eradicated from Europe, grey wolves (*Canis lupus*) have recently undergone a remarkable demographic recovery, but their long-term survival remains precarious. By analyzing 1,001 genomes, we uncover a mosaic of distinct evolutionary lineages, not a single recovering population. Northern populations exhibit Asian wolf ancestry, while southern populations preserve ancient Holocene lineages, with dog introgression varying by region. Isolated wolves from the Scandinavian, Italian, and Iberian peninsulas harbored particularly high levels of inbreeding and fixed deleterious mutations. Signatures of genome erosion were widespread across Europe, with many populations falling well short of the minimum size recommended for long-term survival. Our study reveals the complex tapestry of wolf ancestry and variation across Europe, which calls for nuanced, regional conservation plans founded in genetic monitoring.

## Main Text

Grey wolves (*Canis lupus*) are apex predators that have played a key role in many ecosystems across Eurasia and North America since the mid-Pleistocene. Wolf numbers have declined dramatically across Europe over the past 200 years (*1*). By the mid-20^th^ century, wolves had gone locally extinct in most of western, central and northern Europe, with severe population declines in the Italian and Iberian peninsulas (*1*). In the latter half of the 20^th^ century, wolves underwent a remarkable recovery in Europe, expanding and recolonizing parts of their former range (*2*). Their census sizes reached over 21,500 in 2022, driven by reduced persecution, legal protection, adaptations to a human-dominated landscape, as well as improved habitat quality and connectivity in some regions that enabled long-distance dispersal (*2*). Such recovery has sparked a vivid debate over both conservation management policies and their protection status (*3, 4*). Recently, the Steering Committee of the Bern Convention and the EU Habitats Directive downgraded the protection status of wolves, allowing for increased hunting (*5, 6*).

Despite their widespread distribution, ecological importance, and contentious management policies, there has been no comprehensive, continent-wide genetic study of contemporary wolves across Europe. Most previous work has focused on isolated populations, consistently showing Italian and Iberian wolves as highly diverged from other regions (*1, 7-10*). These populations, along with Scandinavian wolves, also exhibit signs of genomic erosion (loss of diversity, accumulation of deleterious mutations, and reduced adaptive potential) (*10-13*). Moreover, detected dog introgression in multiple regions (*7, 10, 14, 15*), leads to further concerns of long-term viability in European wolves.

To date, management decisions in Europe have relied on 9 populations following the LCIE (Large Carnivore Initiative for Europe) groupings, based on range, numbers, and trends (*16*). Here, by incorporating a continent-wide, genomic assessment of European wolves, we provide an in-depth understanding of their connectivity, hybridization, genomic erosion, and potential vulnerability. We refined population groupings and explored how recent events shaped their genetic landscape, which has direct implications for conservation management policies.

### Population structure and dispersal dynamics of European wolves

We generated a dataset of 940 wolf whole genomes with dense spatial sampling (33 countries across Eurasia), combined with 291 published genomes (Fig. 1A, Table S1). We also included 46 wolf-dog hybrids (40 newly sequenced), and 123 outgroup canids (76 newly sequenced) in our dataset (Table S1, Supplementary Methods). Given the differences in coverage (Table S1), we phased and imputed the genomes using a reference panel of 1,715 canids (*17*). After quality control filtering, we used a final dataset of 1,001 modern Eurasian wolves (650 unrelated) and 52,331,758 biallelic variants for further analysis (Table S2).

**Figure 1:**
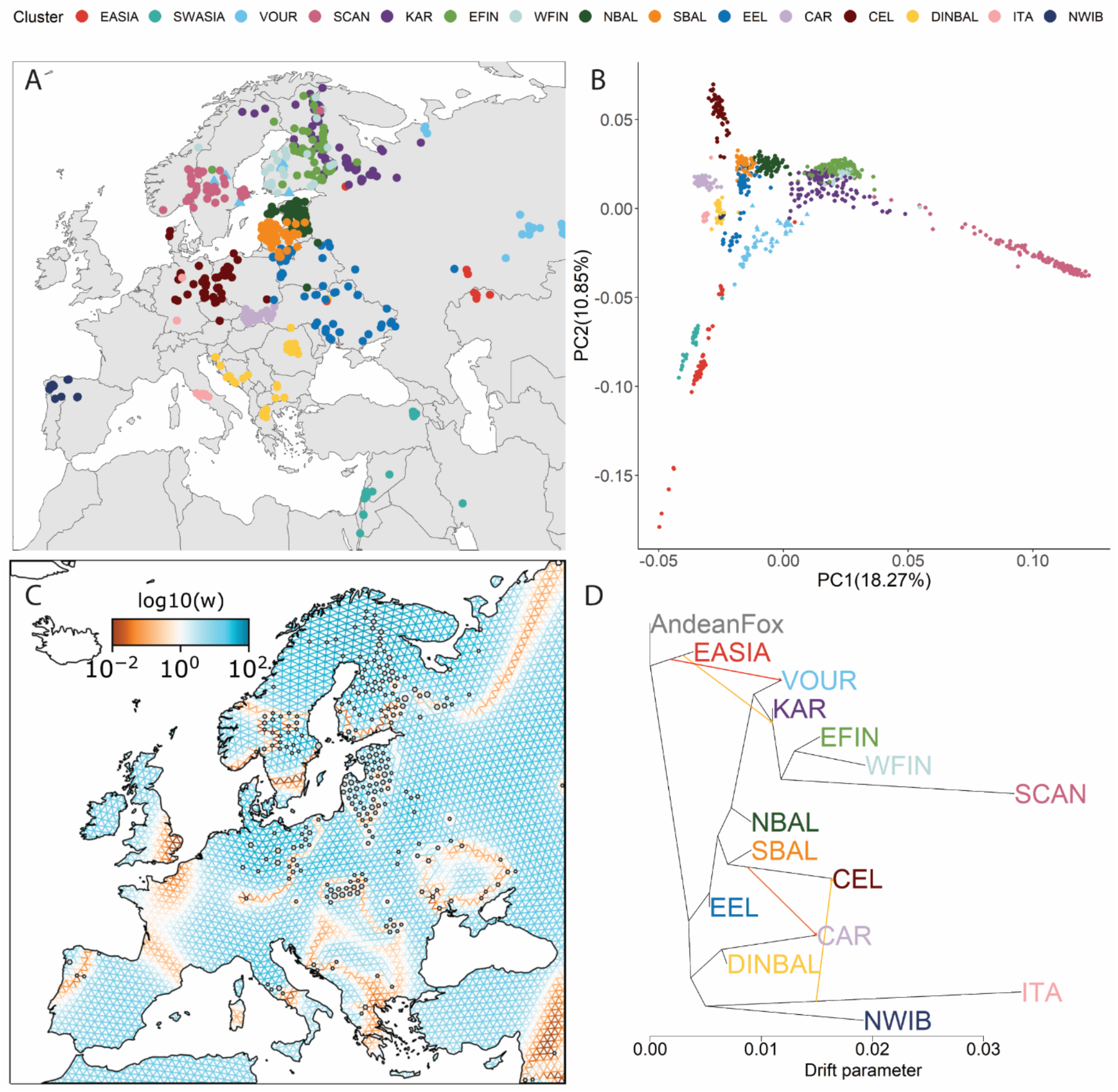
The population structure and dispersal dynamics of contemporary European wolves. **A)** Geographic distribution and Haplonet clustering of unrelated Eurasian wolves (*n*=650) (*18*). **B)** Smartpca of Eurasian wolves (*n*=1,001) colored according to Haplonet clustering (*19*). PC=Principal Component. **C)** Spatial modelling of connectivity amongst Eurasian wolves (*n*=638) using FEEMS (*24*). Sample sizes are reflected by the size of the grey dots at each node. The grid corresponds to a cell area of approximately. 1,550 km^2^ and a cell spacing of 55 km. **D)** Orientagraph topology of wolf clusters (*n*=619) with four migration edges (*23*). SWASIA wolves were excluded due to confounding the topology. Clusters are named according to their main region: EASIA: Eastern Asian, SWASIA: Southwestern Asian, VOUR: Volga-Ural, SCAN: Scandinavian, KAR: Karelian, WFIN: Western Finland, EFIN: Eastern Finland, NBAL: Northern Baltic, SBAL: Southern Baltic, EEL: Eastern European Lowlands, CAR: Carpathian, CEL: Central European Lowlands, DINBAL: Dinaric Balkan, ITA: Italian Peninsula, NWIB: Northwestern Iberian.

To resolve the population structure of wolves, we conducted haplotype-based clustering which identified 15 clusters across Eurasia, broadly corresponding to geography (Figs. 1A, S1-4) (*16, 18*), which were used for grouping in downstream analyses. These clusters were named following LCIE groupings (*16*), although our analysis revealed a more fine-scaled population structure. Principal Component Analysis (PCA) (*19*) also revealed population structure in Eurasian wolves, where the Eastern Asian (EASIA: China, Kazakhstan, Korea, Mongolia, and Russia) and Southwestern Asian (SWASIA: India, Iran, Israel, Türkiye, Saudi Arabia, and Syria) wolves were separated from those in Europe (Figs. 1B, S5). Scandinavian (SCAN: southern Sweden and Norway) and Central European Lowlands (CEL: Austria, Czech-Republic, Denmark, Germany, and Poland) wolves formed clusters distinct from other European individuals, likely reflecting genetic drift from small founder populations (*20, 21*).

Despite geographic structure, our analyses revealed evidence of long-range movements. We found a single cluster of wolves in the Italian Peninsula (ITA), in contrast to the LCIE grouping of separate Apennine and Alpine populations (Fig. 1A) (*16*). Wolves from this cluster were detected as far north as Germany, providing evidence of a northward expansion of this lineage previously restricted to central and southern Apennines (Fig. 1A) (*22*). An admixture graph (Figs. 1D, S7-8) (*23*) modeled the Central European Lowlands cluster as deriving from a Southern Baltic source (SBAL: Latvia and Lithuania), also showing gene-flow with Italian and Carpathian wolves (CAR: Germany, Slovakia, and Poland). This reflects the recent dispersal of wolves from Poland into Germany after their extirpation in the 1850’s, and the subsequent expansion of this new population into surrounding areas(*21*).

However, we still found evidence of reduced gene-flow in the past. On the genetic connectivity map (*24*), Italian wolves were isolated from the Dinaric-Balkan (DINBAL: Bosnia and Herzegovina, Bulgaria, Croatia, Greece, Romania, and Ukraine), Northwestern Iberian (NWIB: Portugal and Spain) and Central European Lowlands clusters (Figs. 1C, S6). Also, Carpathian wolves formed a tight and distinct cluster with reduced gene-flow from surrounding regions, possibly driven by habitat fragmentation, low population numbers and ecological differences (Figs. 1A, 1C) (*25-27*).

Some genetic clusters had different geographic ranges than LCIE, with Romanian wolves grouping with the Dinaric-Balkan cluster rather than Carpathians. We also found distinct genetic clusters of wolves from the Southern Baltic, Northern Baltic (NBAL, from Belarus, Estonia, Latvia, and Lithuania), and Eastern European Lowlands (EEL, from Belarus, Lativa, Lithuania, Poland, Russia, and Ukraine) regions, despite these being considered a single LCIE population (Baltic) (Fig. 1A) (*16*).

The admixture graph showed that wolves from Fennoscandia (Sweden, Norway, Finland, Karelian Russia) originated from the Volga-Ural cluster (Fig. 1D). They subsequently differentiated into four genetic clusters (KAR WFIN, EFIN, SCAN) (Fig. 1B), with the Scandinavian wolves being highly diverged (Figs. 1A, 1D, S4). This is consistent with the known recolonization of Norway and Sweden by an initial breeding pair dispersing from Karelia in 1983, followed by three more migrants (*28*). The barriers detected in the central Scandinavian peninsula (Fig. 1C) can be attributed to reindeer management areas in Finland which hinder wolf dispersal into this area (*29*). Wolves in Finland and Russian Karelia formed three distinct groups, revealing a more fine-grained clustering than defined by LCIE, which classifies all individuals in this region as Karelian. Individuals assigned to the Eastern Asian cluster occurred as far west as Russian Karelia and Ukraine, suggesting that continuous gene-flow has bridged wolves across Asia and eastern Europe (Fig. 1A). Wolves from zoos in Finland, Sweden, and Norway had distinct ancestry from their wild counterparts, clustering with Volga-Ural individuals. Although records are scarce, these individuals may be descendants of Russian wolves (Supplementary Text).

### Different ancestral histories for northern and southern European wolves

Consistent with the westward dispersal of Asian wolves, D-statistics (which measure allele sharing to detect gene flow) showed an excess of Asian-like ancestry in northern and eastern European populations (Figs. 2A, S9-10) (*30*). Excess Eastern Asian ancestry was found only in Fennoscandian and Northern Baltic wolves, likely reflecting recolonization of these regions following population collapses in the mid-20^th^ century (*28*). In contrast, Dinaric-Balkan wolves exhibited limited shared ancestry with Southwestern Asian wolves, likely due to gene-flow barriers (Figs. 1C, S9).

**Figure 2:**
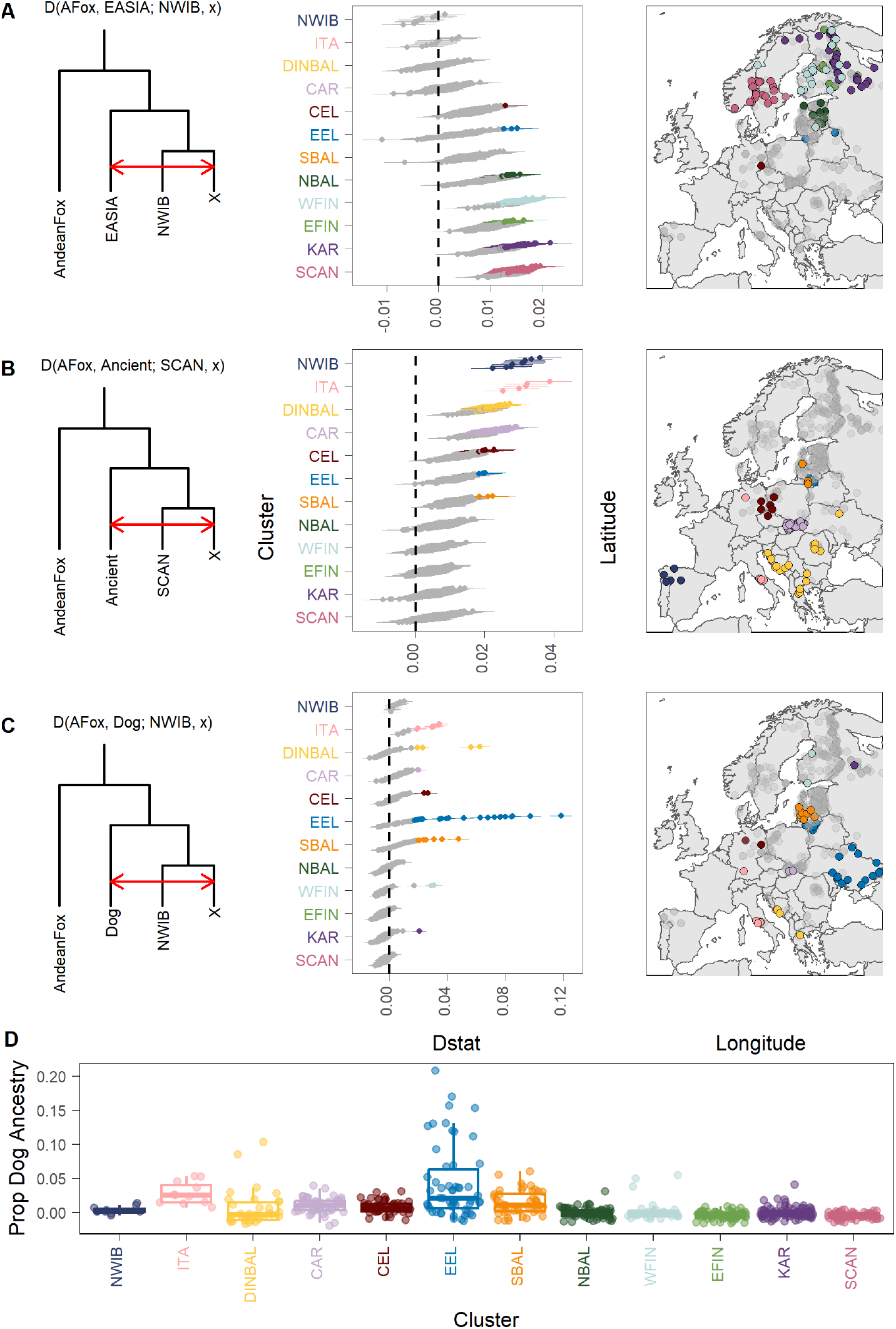
Genetic ancestry and introgression in European wolves. **(A–C)** *D*-statistics analyses (*30*): the left panel shows the topology illustrating significantly positive *D*-statistics; the middle panel presents *D*-statistic estimates for individuals grouped into clusters as defined in Figure 1B; colored points indicate significant positive values (|*Z*| ≥3). the right panel maps the geographic locations of samples showing significant introgression signals (***A:*** Eastern Asian; ***B:*** Ancient Holocene (HMNH-007); ***C:*** Dog). **D)** *f*_*4*_-ratio tests estimating the proportion of dog ancestry in each wolf genome (*30*).

Conversely, wolves from southern Europe, particularly from Italian and Iberian peninsulas, showed negligible genetic affinity to Asian wolves, but strong allele sharing to European wolves from earlier in the Holocene, including a wolf from Hungary dated to 3,500 years ago (*31*) (Fig. 2B, Table S3). This Holocene-like ancestry was also present in the Carpathian and Dinaric-Balkan clusters, which have experienced limited gene-flow from Asia, indicating that these southern regions served as a refugia for ancient ancestral variation (*9*). Contemporary individuals showed no detectable ancestry from any Eurasian Pleistocene wolves (20,000-13,000 years ago) but did show an affinity with Holocene wolves from 10,000 years ago (Table S3), corroborating a previously demonstrated genetic turnover at the Pleistocene/Holocene transition (*32*).

However, no wolves showed excess genetic sharing with an ancient Holocene wolf from Sweden (Table S3), indicating the wolves that inhabited Fennoscandia 5,000 years ago have been replaced (Figs. S11-13).

### Regional variation in dog introgression

Recent population declines in European wolves has raised concerns about increased dog introgression and consequently, genetic swamping, which threatens the genetic integrity, and viability of this species (*4*). Using an *f* 4-ratio test (*30*), we quantified genetic material from European dogs distributed in wolves across Europe. In particular, 47% of the Italian and 46% of the Eastern European Lowlands clusters showed significant dog introgression, as previously described (*7, 14*) (Figs. 2C-D, S14-15, Table S4). In the Southern Baltic and Dinaric-Balkan clusters, 29% and 14% of individuals exhibited dog introgression, respectively (*15*) (Figs. 2C-D, Table S4). Not only do half of the wolves from the Eastern European Lowlands show dog introgression, but they also had notably high levels of dog-like ancestry (up to 21% of their genome), indicating hybridization as recently as two generations ago (Fig. 2D).

However, limited introgression was detected in Fennoscandia and the Central European Lowlands, despite recent bottlenecks, where visually identified F1 hybrids are actively removed from these regions following monitoring of the wild populations (Fig. S16). Our results suggest that higher levels of dog introgression tend to occur in regions with more free ranging dogs and no immediate removal of hybrids, rather than being associated with low numbers of wolves. Finally, D-statistics showed no evidence of introgression from wild canids into European wolves, including golden jackals, which have an expanding and overlapping habitat range (Fig. S17) (*33*).

### Genomic erosion reflects recent demography

The effective population size (*N*_*e*_) of wolves dropped at an average rate of ∼3.5% per generation across all European regions between ∼1800 until 1975 (50 to 10 generations ago) (Figs. 3A, S18-20, Table S5). All clusters, except the Dinaric-Balkan and the Eastern European Lowlands, reached a minimum *N*_*e*_ of > 500 individuals - considered a critical threshold to maintain genetic diversity and evolutionary potential by the Convention on Biological Diversity (*34, 35*). In the last 10 generations, population sizes have remained constant or begun to increase (Figs. 3A, S19), coinciding with improved legal protection and reduced hunting pressures (*1-3*). Nevertheless, 6 out of 15 clusters still fell under the minimum viability threshold (Table S5).

**Figure 3:**
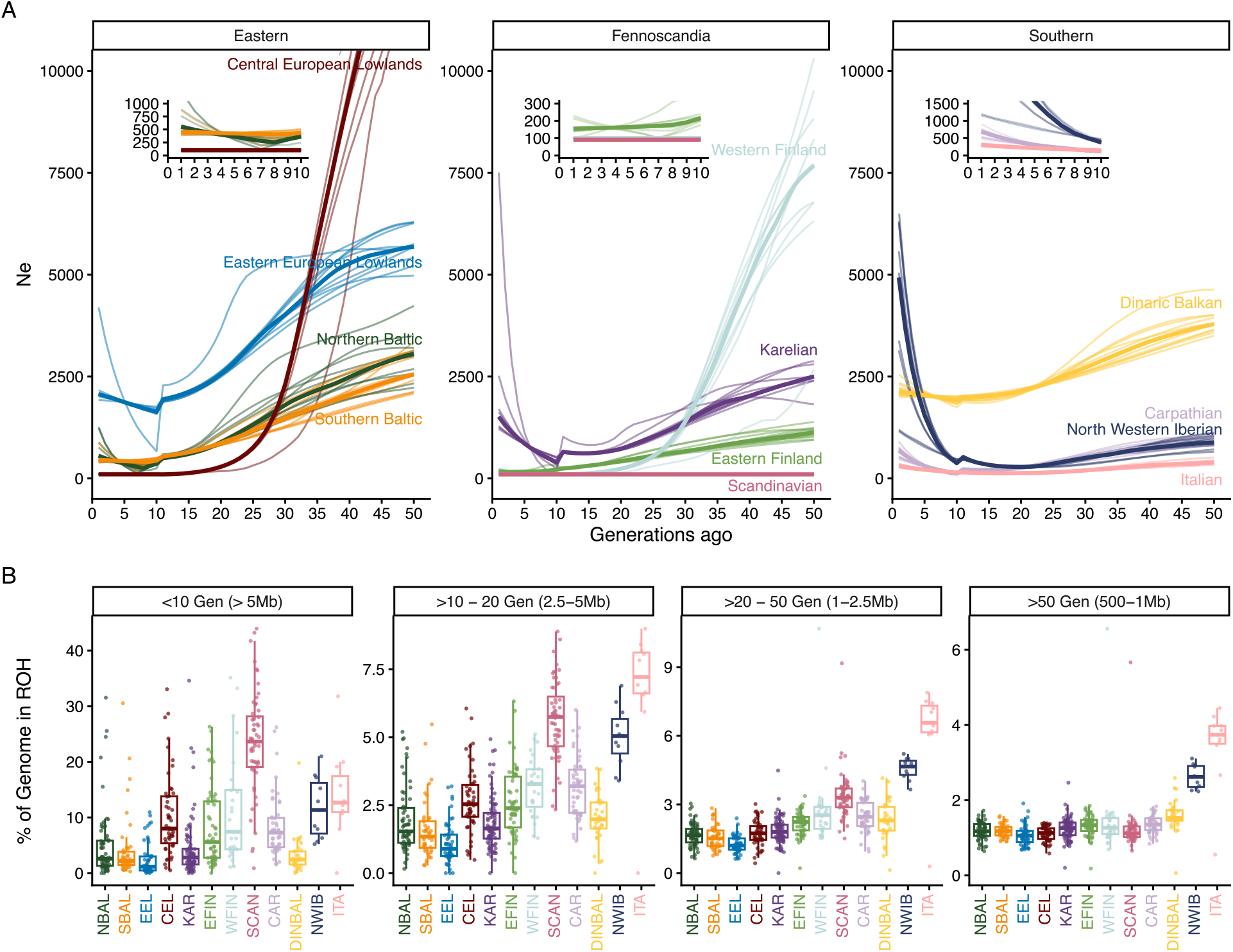
Recent demographic history in European wolves. **A)** Effective population size trajectories in the last 50 generations from HapNe-LD (*42*). The inserts within each facet represent a zoom into the last 10 generations. Thick colored lines represent the median Ne trajectories from 10 independent runs. The light-shaded lines correspond to each independent run. Scandinavia and Karelian3 demographic trajectories overlap in the last 15 generations. **B)** Percentage of the genome in ROH by age (in generations) and sizes (in Mb). ROH lengths (cM) are converted to generations with the formula G = 100/(2 × cM) (*43*), using a 1 cM/Mb recombination rate. The time span of <10 generations correspond to the last 45 years (1975-2020), assuming a generation time of 4.5 years (*44*). The time span of 10 to 20 generations overlaps between 90 and 45 years ago (1930-1975), from 20 to 50 generations overlaps primarily with the 19th century (1795-1930) and >50 generations correspond to the oldest events, before the 19^th^ century.

The differences in demographic trends across clusters, including historical *N*_*e*_, bottleneck strength and population connectivity affects genomic erosion and calls for region-specific conservation management (*11*).

#### Recolonization bottlenecks and diversity in Fennoscandia

Compared to other European wolves, the Scandinavian cluster showed the highest levels of inbreeding (average F_ROH=_0.337), reduced heterozygosity and diversity (Figs. 3B, S21-22). On average, 23.5% of their genome was in long runs of homozygosity (ROH > 5Mb) consistent with their recent founding (*12*). On the other hand, across Fennoscandian wolves, only 1.3% of the genome was found in short ROH (between 500Kb-1Mb) (Fig. 3B), pointing to limited inbreeding in the past, likely because these wolves originated from a larger, interconnected, and diverse source (Figs. 1D, S21). Despite originating from the same source, inbreeding levels and demographic trajectories differed among the Fennoscandian wolves (Figs. 3, S19-20).While the Karelian cluster maintained higher *N*_*e*_ due to ongoing gene-flow from the east (Figs. 2A, 3A), the Scandinavian, Western Finland and Eastern Finland clusters showed extremely low effective population sizes (Fig. 3A, Table S5), likely due to geographic isolation after recolonization. Increases in numbers in these regions has not translated to higher *N*_*e*_ due to the lingering effect of the recent founding bottleneck (*36*).

Fennoscandian wolves showed higher genetic load (∼1% more high-impact and ∼2% more moderate-impact deleterious alleles) than other European population (Fig. 4A, Table S6). This is most notable in Scandinavian wolves, which harbored the highest number of moderate deleterious variants (4% more than the average; Figs. 4A, S23-24, Table S6). Although genetic drift reduced the number of segregating deleterious sites in Scandinavian wolves (∼33% less than the average in European wolves, Figs. 4B, S25, Table S7A), high inbreeding has transformed load into homozygous state, potentially resulting in expression of their deleterious effects (Figs. 4C, S26). In addition, these wolves harbored 1.57 and 2.67 times the average number of fixed high and moderate impact deleterious variants, respectively, more than any other Fennoscandian wolf cluster, indicating reduced efficiency of purifying selection (Fig. 4D, Table S7B).

**Figure 4:**
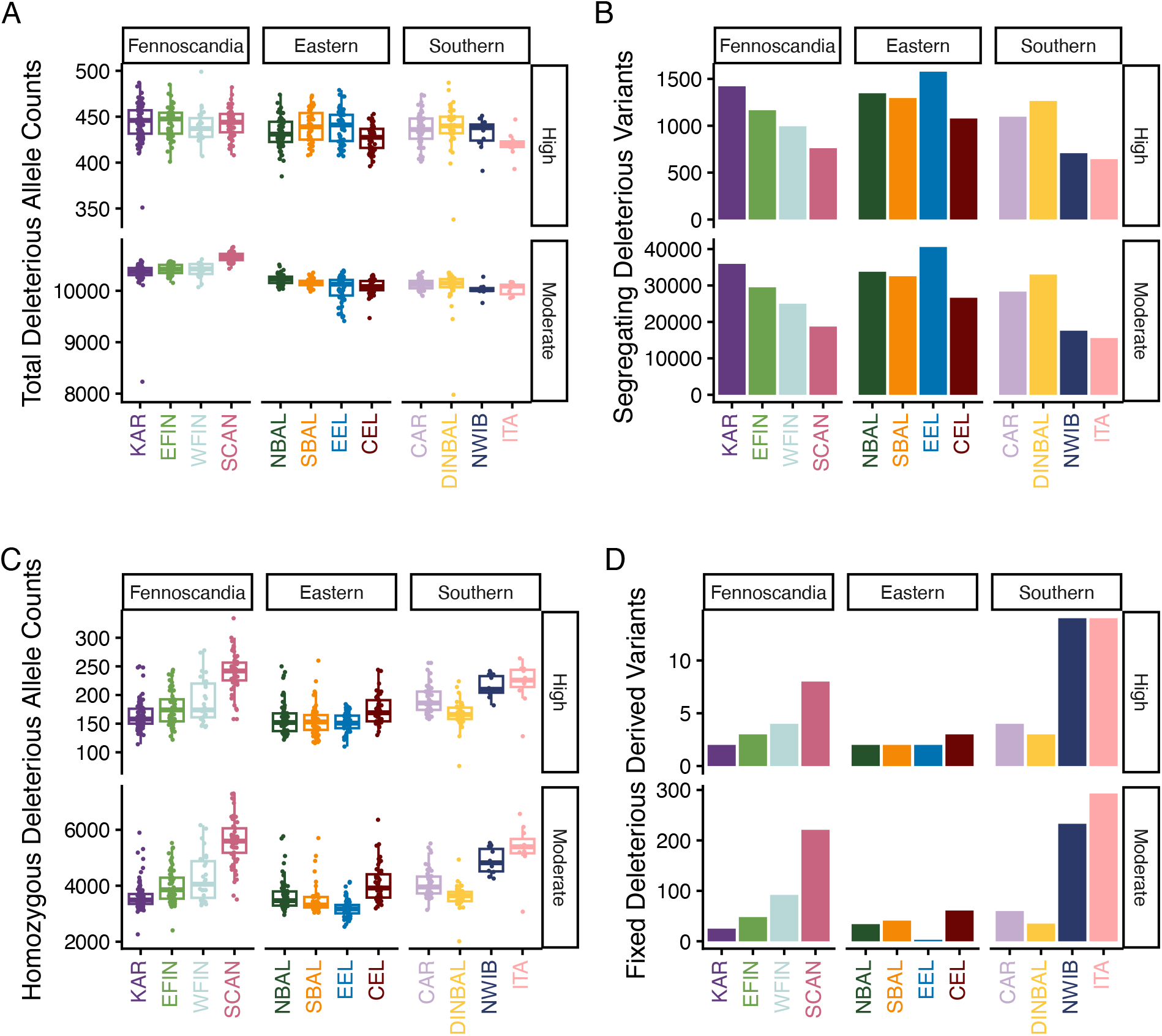
Genetic health in European wolves. **A**) Total deleterious allele counts (moderate and high SnpEff (*45*) categories), which include heterozygous and homozygous allele counts. **B)** Number of segregating deleterious variants per cluster (with frequency between 0 and 1). **C)** Homozygous allele counts for high and moderate categories. **D)** Number of fixed deleterious derived variants per cluster (frequency=1).

#### Long-term isolation consequences in the Italian and Northwestern Iberian wolves

Italian wolves showed the lowest average heterozygosity, closely followed by Scandinavian and Northwestern Iberian wolves (Figs. S21-22, Table S2), and contained high proportions of their genomes in ROHs (F_ROH-ITA_=0.305 and F_ROH-NWIB_=0.239, Figs. 3B, S21, S27). However, unlike Scandinavian wolves, both Italian and Northwestern Iberian wolves exhibited high levels of both long (average 14% of the genome in ROH>5Mb) and short ROH (average 3.5% of the genome in ROH between 500Kb-1Mb) (Fig. 3B), consistent with long-term isolation on their respective peninsulas, coupled with low *N*_*e*_ and inbreeding (Figs. 3, S21, Table S5) (*10*).

Italian and Northwestern Iberian wolves also had the lowest overall number of segregating deleterious variants (∼40% below average) (Figs. 4B, S25, Table S7). Italian wolves, and to a lower degree Northwestern Iberian wolves, carried the lowest genetic load per individual, especially in the highly deleterious category (2% less for moderate-impact; and 4.2% less for high-impact variants than average) (Fig 4A, Table S6). The reduced load is likely the result of genetic drift and purging in populations with persistently low *N*_*e*_ (Figs. 3A, S20). However, the expressed load in homozygous state was higher in both populations, especially in Italian wolves (Figs. 4C, S26). Consistent with long-term isolation, both populations carried the highest numbers of fixed deleterious variants (14 high-impact deleterious variants, 2.75 times the average) (Fig. 4D, Table S7), which could be eased by increased connectivity.

Despite historically low *N*_*e*_, Italian and Northwestern Iberian wolves showed demographic recovery since the 1970s (Figs. 3A, S19-20), consistent with the implementation of conservation measures (*1*). However, we caution that population structure in Northwestern Iberia (*13, 37*) and dog hybridization in Italian wolves could have contributed to the estimated recovery in *N*_*e*_.

#### Recent expansions into the Central European Lowlands

Wolves from the Central European Lowlands showed the strongest decline in *N*_*e*_ (13% decline per generation, Figs. 3A, S19-20, Table S5), reflecting a founder effect resulting from recolonization after 150 years of absence (*3, 21*). Despite this, they exhibited lower recent inbreeding than Scandinavian wolves (10.3% of the genome in long ROH of >5Mb, Figs. 3B, S21), possibly driven by higher connectivity with surrounding clusters (Italian and Carpathian) (Fig. 1C). The combination of recent admixture after the strong founder effect explains the intermediate values of segregating deleterious variants and low number of fixed deleterious variants (3 high-impact deleterious variants) (Figs. 3, S25, Table S7), despite substantial inbreeding (F_ROH_= 0.159, Fig. S21).

#### Reservoirs of genetic variation in Europe at risk

Even wolves from regions in eastern Europe, which maintained high census numbers throughout the 20^th^ century (*1, 2*), showed declines in *N*_*e*_ (at an average rate of ∼3% per generation; Figs. 3A, S19-20, Table S5). The highest current *N*_*e*_ and unsurprisingly, the lowest inbreeding and highest heterozygosity, was found in the Eastern European Lowlands (*N*_*e*_ =2,070), and Dinaric-Balkan clusters (*N*_*e*_ =2,135) (Figs. 3, S21, Table S2). Although higher *N*_*e*_ has masked most of the genetic load in these clusters in a heterozygous state (Fig. 4C, S23, S26), they carried a higher number of segregating deleterious variants (Fig. 4B, 20-40% more than average). This high number of segregating deleterious variants poses a risk in the event of a future population collapse. Furthermore, these clusters exhibited higher proportions of dog admixture (Fig. 2D), which may influence genetic diversity estimates, showing that genetic health measures should be interpreted with caution where interbreeding with dogs occurs (Fig. S28).

### Conclusions and future directions

This study provides the first continent-wide genomic assessment of contemporary European wolves, resolving population structure, dispersal patterns, and connectivity. We improve on the geographic-based LCIE grouping, identifying 15 distinct genetic clusters across Europe, including fine-scale structure in the Baltic, Finnish, and Karelian regions. Northern and central European wolves carry eastern Asian ancestry, whereas southern regions serve as a refugia for ancient Holocene lineages. These patterns reflect the combined legacy of historical extirpation, recolonization, and dispersal patterns, demonstrating that wolves from each region have a unique evolutionary history. Expanding genomic surveys in western Europe, particularly in the Alpine and Pyrenean regions, will further refine our understanding of recent demographic expansions.

Despite an increase in census numbers over the past four decades, many populations remain below the minimum effective population size (*N*_*e*_ > 500) needed to maintain adaptive potential (*34*). Scandinavian, Italian, and Northwestern Iberian wolves harbor 2-3 times more fixed deleterious variants than the average, which has been reported as a strong predictor for extinction risk (*38*). Moreover, inbreeding and loss of genetic variation reveals the long-term consequences of recent population bottlenecks in these wolves, as well as within the newly formed Central European Lowlands population. Hybridization with domestic dogs further threatens genetic integrity in the Eastern European Lowlands, Italian, Southern Baltic, and Dinaric-Balkan wolves, demonstrating hidden genomic risks, even in seemingly large populations.

Our findings provide concrete guidance for conservation management. Connectivity between regions with low diversity, including Northwestern Iberian, Italian, and Central European Lowlands wolves should be encouraged to reduce genomic load and inbreeding (*39*). Encouraging further gene-flow between Asian and Fennoscandian wolves could mitigate genomic erosion, especially in the isolated Western Finland and Scandinavian clusters. However, the benefits of improved connectivity on long-term viability should be further investigated (*40*). At the same time, targeted efforts to limit dog hybridization, particularly in the Eastern European Lowlands are critical to prevent genetic swamping and preserve genomic integrity. The potential impact of newly erected border fences on connectivity should also be closely monitored (*41*).

By integrating fine-scale genomics with demographic insights, we show that each European wolf population has its own unique demographic history. We emphasize that effective conservation should be region-specific and guided by genomic evidence, accounting for differences in demographic histories, connectivity, hybridization, and genomic erosion.

## Acknowledgments

We would like to thank the Association for Nature “Wolf” (Poland) for providing samples. We would also like to thank Kesem Kazes, Bogumiła Jędrzejewska, Włodzimierz Jędrzejewski, Peep Männil, Janis Ozolins, Kamila Plis, Vadim E. Sidorovich, Pedro Silva, Steve Smith, and Rosie Drinkwater for help with sample collection and processing. We would like to thank Cecilie Grønlund Clausen for assistance in the lab.

## Funding

Norwegian Environment Agency (project 18088069) (MTPG, HKS, MDM, XS, SG, MHSS).

ERC Consolidator Award 681396-Extinction Genomics (MTPG).

DNRF143 Center for Evolutionary Hologenomics (MTPG, SG).

National Science Centre, Poland (grant No. 2024/55/B/NZ9/02690) (SN).

Taylor Family-Asia Foundation Endowed Chair in Ecology and Conservation Biology (Grant number 31914.100.004) (BBoldgiv).

Technology Agency of the Czech Republic (project SS07010447) (BČB, PH).

The State Assignment of ZIN RAS (grant 125012800908-0) (GFB, MS).

Institute of Biology Karelian Research Centre of the Russian Academy of Sciences supported by the project FMEN-2022-0003 (KT, AKuznetsova).

The Swedish Research Council award 2021-00625 (LD).

ERC Advanced Grant Award 101054984-PrimiGenomes (LD).

The Knut and Alice Wallenberg Foundation Award 2022.0033 (LD).

For data from Croatia: Bernd Thies Foundation, UK Wolf Conservation Trust, EU LIFE Program, Croatian fund for Nature and Environment, and Plitvice Lakes National Park (JK).

For data from Türkiye: Foundation Segre, Sigrid Rausing Trust, National Geographic Society, STGM, TANAP, TUBITAK (under the 222 N122 Biodiversa+2021 call), Barbara Watkins, BTC Co., and the Whitley Fund (JK).

Portuguese Foundation for Science and Technology (2022.07926.CEECIND) (RG).

Estonian Ministry of Education and Research (grants PRG1209 and TK215) (US).

European Union’s Horizon 2022 research and innovation program under the Marie Sklodowska Curie grant agreements: 101105854 –FENRIR (ETT), 101107083 – ResQ (CF).

Views and opinions are those of the authors only and do not necessarily reflect those of the European Union, the European Research Council or the Research Executive Agency. Neither the European Union nor the granting authority can be held responsible for them.

## Author contributions

Conceptualization: MTPG, SG, M-HSS, MDM, HKS, ETT, CF, XS Funding acquisition: MTPG, SG, M-HSS, MDM, HKS

Samples: MMC, ZA-L, AÅ, LBalčiauskas, LBaltrūnaitė, GFB, BBoldbaatar, BBoldgiv, BČB, TB, DB, PC, AKÇ, EÇ, ME, ØF, LF, EG, JH, LJ, DCK, AAK, ÇK-A, IK, AKoptaz, PK, JK, AKuznetsova, LK, LLarsson, GL, AL, FL, LLlaneza, SLFM, SM, ØØ, LP, MSablin, MSkage, CS, DS, ÇHŞ, RŠ, KT, LT, CV, NY, MÅ, RWM, PH, CN, MN, SN, LD, RG, US, LF, JA, HKS, M-HSS

Lab work: CHS-O, LTL, NFGM, MMC, SSTM

Data analysis: ETT, CF, XS, JRM, GH-A, SG with input from LH. Writing – original draft: ETT, CF, XS

Writing – review & editing: ETT, CF, SG, with input and approval from all co-authors.

## Data and materials availability

Raw sequencing reads for all individuals sequenced in this study have been deposited in the European Nucleotide Archive (accession PRJEB107090). The accession number and metadata for each sample can be found in Table S1.

Scripts and code for population genetic analysis can be found at https://github.com/claudefa/EuropeanWolves_genomics and https://github.com/evelyn-bio/modern_wolves

## Supplementary Materials

Materials and Methods

Figs. S1 to S28

Tables S1 to S27

References (46–*94*)

## References

1. M. Hindrikson et al., Wolf population genetics in Europe: a systematic review, meta-analysis and suggestions for conservation and management. Biol. Rev. 92, 1601–1629 (2017).

2. C. Di Bernardi et al., Continuing recovery of wolves in Europe. PLOS sustain. transform. 4, e0000158 (2025).

3. G. Chapron et al., Recovery of large carnivores in Europe’s modern human-dominated landscapes. Science 346, 1517–1519 (2014).

4. J. C. Blanco, K. Sundseth, The situation of the wolf (Canis lupus) in the European union – An in-depth analysis. A report of the N2K Group for DG Environment, European Commission. (2023).

5. Directive (EU) 2025/1237 of 17 June 2025, amending Council Directive 92/43/EEC as regards the protection status of the wolf (Canis lupus) (2025) OJ L, 2025/1237

6. Proposal (EU) of 27 September 2024, Proposal to amend Appendices II and III of the Bern Convention of the Conservation of European Wildlife and Natural Habitats by moving the wolf (Canis lupus) from Appendix II to Appendix III (2024) T-PVS/Inf(2024)15

7. A. V. Stronen et al., North-south differentiation and a region of high diversity in European wolves (Canis lupus). PLOS ONE 8, e76454 (2013).

8. M. Pilot et al., Genetic variability of the grey wolf Canis lupus in the Caucasus in comparison with Europe and the Middle East: distinct or intermediary population? PLOS ONE 9, e93828 (2014).

9. P. Silva et al., Genomic evidence for the old divergence of Southern European wolf populations. Proc. Biol. Sci. 287, 20201206 (2020).

10. D. Battilani et al., Beyond population size: whole-genome data reveal bottleneck legacies in the peninsular Italian wolf. J. Hered. 116, 10–23 (2025).

11. C. van Oosterhout et al., Genomic erosion in the assessment of species’ extinction risk and recovery potential. J. Hered., esag011 (2026).

12. M. Kardos et al., Genomic consequences of intensive inbreeding in an isolated wolf population. Nat. Ecol. Evol. 2, 124–131 (2018).

13. I. Salado et al., Large variance in inbreeding within the Iberian wolf population. J. Hered. 115, 349–359 (2023).

14. R. Lorenzini, A. Pizzarelli, L. Attili, M. Biagetti, C. Sebastiani, P. Ciucci, Genetic evidence reveals extensive wolf-dog hybridisation in peninsular Italy: warnings against ineffective management. Biol. Conserv. 313, 111615 (2026).

15. J. Kusak et al., Wolf-dog hybridization in Croatia. Vet. Arh. 88, 375–395 (2018).

16. P. Kaczensky et al., Large carnivore distribution maps and population updates 2017–2022/23. Report to the European Comission under contract N° 09.0201/2023/907799/SER/ENV.D.3 “Support for Coexistence with Large Carnivores”, “B.4 Update of the distribution maps”. IUCN/SSC Large Carnivore Initiative for Europe (LCIE) and Istituto di Ecologia Applicata (IEA) (2024).

17. K. Bougiouri et al., Imputation of ancient canid genomes reveals inbreeding history over the past 10,000 years. Proc. Natl. Acad. Sci. U.S.A. 122, e2416980122 (2025).

18. J. Meisner, A. Albrechtsen, Haplotype and population structure inference using neural networks in whole-genome sequencing data. Genome Res. 32, 1542–1552 (2022).

19. A. L. Price, N. J. Patterson, R. M. Plenge, M. E. Weinblatt, N. A. Shadick, D. Reich, Principal components analysis corrects for stratification in genome-wide association studies. Nat. Genet. 38, 904–909 (2006).

20. M. Åkesson et al., Genetic signature of immigrants and their effect on genetic diversity in the recently established Scandinavian wolf population. Conserv. Genet. 23, 359–373 (2022).

21. A. Jarausch, V. Harms, G. Kluth, I. Reinhardt, C. Nowak, How the west was won: genetic reconstruction of rapid wolf recolonization into Germany’s anthropogenic landscapes. Heredity 127, 92–106 (2021).

22. E. Fabbri et al., From the Apennines to the Alps: colonization genetics of the naturally expanding Italian wolf (Canis lupus) population. Mol. Ecol. 16, 1661–1671 (2007).

23. E. K. Molloy, A. Durvasula, S. Sankararaman, Advancing admixture graph estimation via maximum likelihood network orientation. Bioinformatics 37, i142–i150 (2021).

24. J. Marcus, W. Ha, R. F. Barber, J. Novembre, Fast and flexible estimation of effective migration surfaces. eLife 10, e61927 (2021).

25. W. Jędrzejewski et al., Prey choice and diet of wolves related to ungulate communities and wolf subpopulations in Poland. J. Mammal. 93, 1480–1492 (2012).

26. A. Morales-González, A. Fernández-Gil, M. Quevedo, E. Revilla, Patterns and determinants of dispersal in grey wolves (Canis lupus). Biol. Rev. 97, 466–480 (2022).

27. M. Hindrikson, J. Remm, P. Männil, J. Ozolins, E. Tammeleht, U. Saarma, Spatial genetic analyses reveal cryptic population structure and migration patterns in a continuously harvested grey wolf (Canis lupus) population in north-eastern Europe. PLOS ONE 8, e75765 (2013).

28. M. Åkesson, O. Liberg, H. Sand, P. Wabakken, S. Bensch, Ø. Flagstad, Genetic rescue in a severely inbred wolf population. Mol. Ecol. 25, 4745–4756 (2016).

29. I. Kojola, S. Kaartinen, A. Hakala, S. Heikkinen, H.-M. Voipio, Dispersal behavior and the connectivity between wolf populations in northern Europe. J. Wildl. Manag. 73, 309–313 (2009).

30. R. Maier, P. Flegontov, O. Flegontova, U. Işıldak, P. Changmai, D. Reich, On the limits of fitting complex models of population history to f-statistics. eLife 12, e85492 (2023).

31. A. Bergström et al., Grey wolf genomic history reveals a dual ancestry of dogs. Nature 607, 313–320 (2022).

32. J. Ramos-Madrigal et al., Genomes of Pleistocene Siberian wolves uncover multiple extinct wolf lineages. Curr. Biol. 31, 198-206. e198 (2021).

33. R. Rutkowski et al., A European Concern? Genetic Structure and Expansion of Golden Jackals (Canis aureus) in Europe and the Caucasus. PLOS ONE 10, e0141236 (2015).

34. I. Franklin, in “Evolutionary change in small populations” in Conservation Biology: An Evolutionary-Ecological Perspective. (Sinauer Associates, U.S.A., Sunderland, Massachusetts, 1980), pp. 135–149.

35. S. Hoban et al., Monitoring status and trends in genetic diversity for the Convention on Biological Diversity: An ongoing assessment of genetic indicators in nine countries. Conserv. Lett. 16, e12953 (2023).

36. R. Gargiulo, K. B. Budde, M. Heuertz, Mind the lag: understanding genetic extinction debt for conservation. Trends Ecol. Evol. 40, 228–237 (2025).

37. P. Silva et al., Cryptic population structure reveals low dispersal in Iberian wolves. Sci. Rep. 8, 14108 (2018).

38. N. Dussex, H. E. Morales, C. Grossen, L. Dalén, C. van Oosterhout, Purging and accumulation of genetic load in conservation. Trends Ecol. Evol. 38, 961–969 (2023).

39. M. Kardos et al., The crucial role of genome-wide genetic variation in conservation. Proc. Natl. Acad. Sci. U.S.A. 118, e2104642118 (2021).

40. C. C. Kyriazis, R. K. Wayne, K. E. Lohmueller, Strongly deleterious mutations are a primary determinant of extinction risk due to inbreeding depression. Evol. Lett. 5, 33–47 (2021).

41. K. Nowak et al., Weaponizing Europe’s borders imperils wildlife. Science 389, 358–359 (2025).

42. R. Fournier, Z. Tsangalidou, D. Reich, P. F. Palamara, Haplotype-based inference of recent effective population size in modern and ancient DNA samples. Nat. Commun. 14, 7945 (2023).

43. E. A. Thompson, Identity by descent: Variation in meiosis, across genomes, and in populations. Genetics 194, 301–326 (2013).

44. L. Smeds, I. Kojola, H. Ellegren, The evolutionary history of grey wolf Y chromosomes. Mol. Ecol. 28, 2173–2191 (2019).

45. P. Cingolani et al., A program for annotating and predicting the effects of single nucleotide polymorphisms, SnpEff. Fly 6, 80–92 (2012).

