## Supplementary Materials for "The population structure and genetic health of European wolves"

**The PDF file includes:**

Materials and Methods
Supplementary Text
Figs. S1 to S28
Tables S1 to S7
References 46 to 94

### Materials and Methods

Sample collection and ethics

- **Sample collection**

We collected wolf samples from across Eurasia to capture their broad geographic distribution, alongside suspected wolf-dog hybrids and golden jackals (*Canis aureus,* a wild canid also found in Eurasia). In total, we obtained 1,021 hair, tissue or blood samples for sequencing, consisting of: 905 grey wolves (*Canis lupus*), 20 golden jackals (*Canis aureus*), 56 Norwegian dogs (*Canis lupus familiaris*), and 40 hybrids (*Canis lupus* × *Canis lupus familiaris*).

We supplemented our dataset with 361 publicly available genomes from the European Nucleotide Archive, including: 308 grey wolves (*Canis lupus*), 33 dogs (*Canis lupus familiaris*), five hybrids (*Canis lupus* x *Canis lupus familiaris*), one side-striped jackal (*Canis adustus)*, one golden jackal (*Canis aureus*), four coyotes (*Canis latrans*), two African wolves (*Canis lupaster*), one dingo (*Canis lupus dingo*), one black backed jackal (*Canis mesomelas*), one Ethiopian wolf (*Canis simensis*), one dhole (*Cuon alpinus*), one Andean fox (*Lycalopex culpaeus)*, and two African wild dogs (*Lycaon pictus*). Geographical coordinates were inferred for samples without exact sampling location based on the region specified. Accession numbers for all newly sequenced and publicly available genomes used in this study can be found in Table S1 (*12, 32, 46-64*).

- **Ethics**

Museum specimens, legally hunted animals, necroscopy samples, and postmortem samples from euthanized or roadkill animals did not require ethical approval. The ethics for each individual sample are detailed in Table S1.

The wolf samples from Türkiye were collected by specially trained scientists with all procedures involving animal handling approved by the Republic of Türkiye Ministry of Agriculture and Forestry. The work was conducted under the supervision of the Kafkas University Animal Experiments Local Ethics Committee (Approval No. KAU-HADYEK/2018-050). In addition, the study was conducted under research permits issued by the Ministry of Agriculture and Forestry, General Directorate of Nature Conservation and National Parks (Permit No. 72784983-488.04-151690 and 72784983-488.04-114100).

The wolf samples from Poland were collected with permission from the Directorate for Environmental Protection (permits No. DZP-WG.6401.08.2.2015.JRO and DZP-WG.6401.08.1.2017.bp).

The dog samples from Norway were obtained from the biobank at The Norwegian University of Life Sciences where the owners signed a written consent allowing the DNA to be used for research.

DNA extraction and sequencing

Samples were classified based on preservation conditions into high-quality or degraded DNA. For high-quality samples, DNA was extracted using the KingFisher™ Duo Prime Purification System following the manufacturer's protocol. Libraries were prepared either in-house at the University of Copenhagen or by BGI Copenhagen. For degraded samples, DNA extraction followed protocols optimized for degraded (ancient) DNA (*32, 60, 61, 65, 66*), and libraries were constructed using either BEST (*67*) or SCR (*68*) methods. The libraries were sequenced with the DNBSEQ-G400 platform using the BGI commercial service (Copenhagen, Denmark), either as paired-end sequencing (PE150) or single-end sequencing (SE100).

Sequence alignment, variant calling, and imputation

- **Sequence alignment**

The sequencing reads in each raw FASTQ file were aligned to the CanFam 3.1 reference genome (*69*) using the Paleomix pipeline (v1.2.13.2) (*70*). Reads were demultiplexed, trimmed, and collapsed using AdapterRemoval (v2) (*71*) using the parameters: --mm: 3 --minlength: 25 --qualitymax: 93 --collapse: yes --trimns: yes --trimqualities: yes. Reads were aligned using BWA backtrack (v0.7.15) (*72*) with the parameters: --MinQuality: 0 --FilterUnmappedReads: yes --UseSeed: no. The resulting BAM files were realigned around INDELs using GATK (v3.4) (*73*), and PCR duplicates were removed using Picard Tools (v1.128) (http://broadinstitute.github.io/picard).

- **Variant calling and imputation.**

Genotypes were imputed and phased using GLIMPSE v1.1.1 (*74*) per autosome, with a canid reference panel containing 139,268,526 variants and 1,715 canids, including 116 grey wolves (*17*).

Imputation was performed following the pipeline outlined in the GLIMPSE documentation. First, variants were called from the mapped BAM files using BCFtools (v1.12) (*75*) mpileup with the parameters: “--ignore-RG -I -E -q 30 -Q 20 -a 'FORMAT/DP”, and BCFtools call with the parameters: “-Aim -C alleles”, conditioning on the variants in the reference panel and running each chromosome separately. Quality filtering of variants was performed using BCFtools view with the parameters: “AC<=0 || AN <=100 || QUAL="."”.

For post-imputation quality control, the imputed genotypes were filtered using an INFO score threshold of 0.8 (*76*), resulting in 1,382 individuals and 52,331,758 autosomal, biallelic SNPs for further analysis (Table S1).

Related individuals were identified using KING (v2.3.1) (*77*), and duplicates and historical samples were removed. Pairs with relatedness > 0.177 were considered related; one individual per pair was excluded in analyses requiring unrelated samples (Table S2).

Population genetic analysis

- **Clustering Analysis**

We used the phased and imputed dataset to group the Eurasian wolves into genetic clusters. We removed duplicates, wild canids, dogs, known hybrids, and North American wolves from the dataset, leaving 1,001 Eurasian wolves. To avoid biases from related individuals, we used 650 unrelated Eurasian wolves for further analysis.

We conducted an unsupervised ADMIXTURE analysis of variants with an MAF>0.01 (19,091,689 variants). The variants were linkage disequilibrium (LD) pruned using PLINK (v1.9) (*78*), using the parameter --indep-pairwise 500 50 0.1, leaving 310,241 variants.

ADMIXTURE analyses (v1.3) (*79*) were performed for K values between 3-18 (Fig. S1), with cross validation errors identifying the optimal model as K=15.

We further explored population clustering using the haplotype-based method implemented in Haplonet (v1.0) (*18*), which applies a variational autoencoder to model haplotypes and infer ancestry components. The analysis was run on the same dataset (650 unrelated wolves, 19,091,689 variants), training the model on each chromosome independently. Haplonet admix was run on all chromosomes for K values from 3 to 18 (Fig. S2). We chose a K value of 15 to group the wolves, as it best represented the geographical clustering of the samples (Fig. 1A, S3). Each wolf was assigned to the cluster in which it showed the highest ancestry proportion, and clusters were labeled by their dominant geographic distribution for subsequent analyses, in line with previous grouping names from the LCIE (*16*). Related wolves were assigned to the same cluster as their close relative identified in the KING analysis.

- **Neighbour Joining Tree**

To visualize relationships among Eurasian wolf clusters, we constructed a neighbor-joining (NJ) tree using the 650 unrelated individuals and the 310,241 LD-pruned SNPs. Pairwise distances were calculated using PLINK (v1.9), using the parameter --distance square 1-ibs flat-missing. A BioNJ tree was inferred using FastMe (v2.6.1.3) (*80*), with 100 bootstrap pseudoreplicates. The resulting topology indicated shallow branch lengths among different Eurasian wolf clusters, consistent with a rapid divergence and/or ongoing gene-flow amongst groups (Fig. S4).

- **Principal Components Analysis (PCA)**

We performed principal components analysis (PCA) on the phased and imputed variant dataset to investigate population structure among Eurasian wolves. Variants were filtered for a minor allele frequency (MAF) threshold of 0.01 using BCFtools view (v1.2.0), resulting in 1,001 wolves and 18,150,192 variants. To avoid biases from related individuals, we used the 650 unrelated, Eurasian wolves to estimate the principal components and projected the related individuals using the smartpca function of Eigensoft (v8.0.0) (*19, 81*). The wolves were colored according to their cluster estimated from the Haplonet admix analysis (Fig. 1B, S5).

- **OrientAGraph**

To model population relationships and historical gene-flow among clusters, we used OrientAGraph (v1.2) (*23*), with the LD pruned SNPs ($n$=310,241), and the unrelated Eurasian wolves with the Andean Fox (*Lycalopex culpaeus*) included as an outgroup ($n$=651). We ran ten replicates with migration edges ranging from 0 to 4 (-mlno mode) and identified the optimal number of migration edges using a linear model implemented in OptM (v1.0) (*82*). Because the Southwestern Asia (SWASIA) cluster exhibited excess allele sharing with the Andean fox (Fig. S7), likely reflecting introgression from other wild canids, we reran the OrientAGraph analysis excluding this population to confirm topological stability ($n$=619), where the optimal number of migrations edges was 4 (Fig. 1C, S8).

- **FEEMS**

We ran Fast Estimation of Effective Migration Surfaces (FEEMS) (v1) (*24*) with the LD pruned SNPs ($n$=307,508) and excluding 12 captive wolves (638 wolves; Table S2). For the map including all of Eurasia we used grid_100, which corresponds to res = 6 in dgconstruct from package dggridR (https://github.com/r-barnes/dggridR/). For the map including just Europe we aimed for higher resolution by generating a finer grid, corresponding to res=7 in the dgconstruct library. To run the software we followed the recommendations available from https://github.com/NovembreLab/feems.

- **D-statistics**

We characterized the ancestral composition of modern wolf genomes and detected introgression from other canids using D-statistics with the formula D=(BABA-ABBA)/(BABA+ABBA). Analyses were performed using admixtools2 (v2.0.4) (*30*) on each unrelated Eurasian wolf (*n*= 650). To minimize potential deamination and sequencing error effects for the ancient genomes, we restricted analyses to transversion variants with a MAF ≥ 0.01 (*n*=5,261,623). The Andean fox (*Lycalopex culpaeus*) served as a consistent outgroup to root all comparisons. Across all tests, Z-scores > |3| were considered significant. Directionality and magnitude of the D-statistics were used to infer gene-flow between specific clusters or species.

*Asian introgression*

We tested for gene-flow from Asian into European wolves using D-statistics in the form: D(AndeanFox, Asian Wolf; European Wolf, x). A significantly positive D-statistic indicates introgression of Asian wolf genetic material into European wolves. Asian wolves from each of the three clusters identified in the Haplonet admix analysis were placed in the H2 position: VOUR (MW1063), SWASIA (MW167), and EASIA (MW522). In the H3 position, we used Scandinavian (MW145) and Northwestern Iberian (MW122) wolves, representing the two dominant modern European lineages. All European wolves (*n*=537) were placed in the H4 position (Fig. 2A, S9).

We then tested for gene-flow from different wolves across Asia in the form D(AndeanFox, x, Asian Wolf1, Asian Wolf2), where a positive D-statistic means excess allele sharing with Asian Wolf2, and a negative D-statistic means excess allele sharing with Asian Wolf 1. The wolves tested from across Asia were: MW497 (St. Petersburg), MW1076 (Kazakhstan), MW535 (Yakutsk), Chinese_CAN16 (Xinjiang), Chinese_CAN9 (Tibet), and KoreanWolf4 (Korea) (Fig S10).

*Ancient wolves*

We compared modern wolves to the 14 ancient Eurasian wolves (<18 ka BP, > 0.5X depth of coverage, Table S3) to assess genetic continuity and ancient introgression. Ancient sequences were pseudo-haploidized in ANGSD (v0.940) (*83*), conditioning on the modern transversion sites with the parameters: -dohaplocall 1 -doCounts 1 -doMajorMinor 3 -remove_bads 1 -C 50 -minMapQ 25 -minQ 30 -uniqueOnly 1 -baq 1 -doPlink 2 -doGeno -1. The pseudohaploid variant calls were merged with the modern samples using the mergeit function in the Eigensoft package (v8.0.0) (*19, 81*).

D-statistics were calculated in the form: D(AndeanFox, Ancient Wolf; Modern Wolf, x). Ancient wolves were placed in the H4 position, with Scandinavian (MW145) and Northwestern Iberian (MW122) wolves as H3 (Figs. 2C, S11, S12). To avoid confounding by dog introgression, we excluded wolves with significantly positive values in the form D(AndeanFox, GermanShephard, NWIB, x) (*n*=46), leaving 491 European wolves in the dataset (Fig. S13).

We measured whether the modern wolves were more closely related to dogs or ancient wolves through D-statistics in the form D(AndeanFox, x, Ancient, Dog). A significantly positive D-statistic indicates excess allele sharing with dogs, where a significantly negative D-statistic indicates greater allele sharing with the ancient wolf. All modern wolves were placed in the H2 position (*n*=650), the GShepDog was used in the H3 position, and all 14 of the ancient wolves (Table S3) in the H4 position (Fig, S13).

*Dogs*
We quantified introgression from domestic dogs using D-statistics in the form: D(AndeanFox, Dog; Wolf, x). A significantly positive D-statistic indicates excess allele sharing with dogs. Dogs representing the six major global ancestry clusters following (*84, 85*) were placed in the H2 position: Sled Dog (AlaskanHusky_SY001), African Dog (BasenjiDog), Sahul Dog (Novembre_Dingo), Asian Dog (HebeiDog), Middle Eastern Dog (AfghanDog), and European Dog (GShepDog). Scandinavian (MW145) and Northwestern Iberian (MW122) wolves were again used as H3 representatives of the major modern European lineages (Figs. 2C, S14).

We inferred the source of dog ancestry using D-statistics in the form: D(AndeanFox, x; Dog1, Dog2). The dogs from the six ancestries were placed in combinations of Dog1 and Dog2, where a significantly positive D-statistic indicates excess allele sharing from the Dog2 lineage, and a significantly negative D-statistic suggests allele sharing from the Dog1 lineage (Fig. S15).

*Wild canids*
To assess introgression from wild canids into modern wolves, we calculated D-statistics in the form: D(AndeanFox, Wild Canid; Wolf, x). Significantly positive D-values indicates excess allele sharing between modern wolves and the wild canid, consistent with gene-flow or shared ancestry. We tested wild canids from Africa, Eurasia, and the Americas in the H2 position: African wolf ALG (Algerian_grey_wolf), Dhole (BerlinZoo), Side-striped jackal (C_adustus), Black-backed jackal (C_mesomelas), Coyote (CaliforniaCoyote), Ethiopian Wolf (Ethiopian_grey_wolf), African Wolf KEN (KenyaJackal), Golden Jackal (Syrian_jackal), African wild Dog RSA (Kruger), African Hunting dog ZWE (Zimbabwe1). As in the Asian D-statistics analyses, Scandinavian (MW145) and Northwestern Iberian (MW122) wolves were placed in the H3 position (Fig. S17).

- ***f*4 ratio test**

We quantified the proportion of dog ancestry in modern wolves using an *f*4-ratio test, calculated as: *f*4(FinnishLapphund, AndeanFox; x, Wolf)/*f*4(FinnishLapphund, AndeanFox; GShepDog, Wolf), where x includes all contemporary European wolves (*n*=537), with a Scandinavian wolf (MW145) placed in the H4 position. This approach provides an estimate of the fraction of wolf genomes derived from dog ancestry relative to a reference dog lineage. The *f*4-ratio tests were estimated using admixtools2 (v2.0.4) (*30*) (Fig. 2D, Table S4).

We also estimated the proportion of dog ancestry in 24 known hybrids from Europe and three wolves identified as hybrids from the KING relatedness analysis (MW1139 from Greece and MW701A from Bulgaria) (Fig. S16).

- **Genetic diversity**

We calculated genome-wide nucleotide diversity ($\pi$) for wolves sampled from each European country using VCFtools (v0.1.17) (*86*). All 52,331,758 sites were used, and diversity was calculated for wolves, grouped by country, excluding duplicates. Diversity was computed in sliding windows of 5 kb (--window-pi 5000) and averaged across the genome to obtain per-country estimates (Fig. S22).

We estimated relative heterozygosity per individual (650 unrelated individuals) with vcftools and vcflib (option vcfhetcount) (v1.0.2) (*87*) on the VCF containing 52,331,758 variants (see above). Relative heterozygosity is defined as the number of heterozygote sites by the total number of sites in the VCF (52,331,758 sites) (Fig. S21A).

- **Inbreeding and runs of homozygosity**

We estimated the inbreeding coefficient per individual as the proportion of the genome in runs of homozygosity ($F_{ROH}$). We obtained runs of homozygosity (ROH) with BCFtools roh (v1.20) with the following parameters: -G 30 --estimate-AF –m genetic_map_CHROM_average_canFam3.1.txt, on a subset VCF including 650 unrelated individuals and variants with a MAF>0.01 (19,091,689 sites). We used the average recombination map per chromosome described for dogs (*88*). We kept only ROH that were > 500Kb and with quality >50 (Fig. S21B). We then stratified $F_{ROH}$ by ROH length with the following bins (500Kb-1Mb, 1-2.5Mb, 2.5-5Mb and >5Mb) and estimated the inbreeding age by converting ROH lengths (cM) to generations with the formula $G = 100/(2 \times cM)$ (*43*), using a 1 cM/Mb recombination rate (*55, 88*). The ROH lengths chosen are then related to background inbreeding (> 50 generations ago), older (20-50 generations ago), intermediate (10-20 generations ago) and very recent inbreeding (<10 generations), respectively (*89*) (Fig. 3B).

- **Genetic load**

We annotated putative deleterious variants in the VCF including 829 canid genomes without related individuals (*n*=52,331,758) using SnpEff (v5.3a) (*90*). We used the existing genome annotation for the dog genome (CanFam3.1.99) within the SnpEff database.

From the annotated VCF, we extracted variants classified as high, moderate, and low impact. High-impact variants are assumed to have a high (disruptive) impact on the protein, probably causing protein truncation, loss of function (LoF) or triggering nonsense-mediated decay (i.e., stop codons, splice donor variants and splice acceptor, start codon lost, etc.). Moderate-impact variants are non-disruptive variants that may alter protein effectiveness (i.e., missense variants). Low-impact variants are mostly harmless or unlikely to change protein behavior (i.e., synonymous variants).

We polarized the discovered variants into ancestral and derived using 3 closely related outgroups (Coyote – sampleIDAlabamaCoyote; golden jackal – sampleID MW692; and African golden wolf – sampleID Algerian_grey_wolf). We considered only sites where at least two out of the three outgroups were homozygous reference (0|0), which we defined as the ancestral state. Any alternative allele present in the wolf was then considered derived, which SnpEff assumes to be the potentially deleterious variant. We required a minimum of two consensuses across outgroups to increase the reliability of polarisation by minimizing the misidentification of ancestral alleles due to lineage-specific substitutions. With this filter, we retained 92.58% of the annotated sites (466,503/503,892) and treated the derived allele as putatively deleterious and the reference as ancestral (Fig 4A). As a double check, we also polarized variants, adding two more distant outgroups (Andean fox – sampleID AndeanFox and Dhole – sampleID BerlinZoo), and requiring a minimum of four consensus across outgroups, which retained 85.99% of sites (433,292/503,892), resulting in very similar estimates of genetic load (Fig. S23).

To approximate the genome-wide genetic load, we counted derived alleles relative to the outgroup ancestral alleles as putatively deleterious. We separated the total genetic load into heterozygous and homozygous components by counting the number of derived alleles at low, moderate, and high predicted levels for homozygous alleles (multiplied by two) and for heterozygous alleles. The homozygous counts are a proxy of the realized load (deleterious variants that express fitness effects); heterozygous counts are a proxy of the masked load (deleterious variants whose fitness effects remain largely hidden). Heterozygous variants can partially express their deleterious fitness effects (*91*), but in the absence of dominance coefficients (h) estimates, we assumed that h ~0 for moderately and highly deleterious mutations (*92*) (Fig 4A, 4C, S26). Next, we estimated allele frequencies per population and calculated Rxy (*93*) per X population using Eastern Asian as Y, since they show to have the largest and most stable $N_e$ trajectory (Fig. S24).
Rxy was calculated as follows: $Rxy = \frac{F_{pop} \cdot (1 - F_{EASIA})}{F_{EASIA} \cdot (1 - F_{pop})}$
(*93*), where $F_{pop}$ represents the allele frequency in each population, and $F_{EASIA}$ represents the allele frequency in the Eastern Asian population. The Rxy values for both high and moderate categories were then normalized with the low Rxy value. We performed a jackknife analysis to estimate the variance around the Rxy values by excluding 1% sites each time, for 100 times, for moderate impact variants, and 10% of the sites, for 10 times, for the high impact variants, and calculated the corresponding Rxy values.
We also counted the number of segregating and fixed derived deleterious variants (for Moderate and High-impact classifications) in each cluster by filtering variants with frequency>0 and <1 for segregating and frequency=1 for fixed (Fig 4B, 4D, S25).

- **Recent demographic trajectories**

We estimated changes in recent effective population size ($N_e$) using HapNe-LD (*42*) over the last 50 generations (the period from 1795 to 2020). We first split the VCF, including 650 unrelated samples with 19,091,689 sites per chromosome and per group, with BCFtools (v1.20) and then converted it to map and ped files with PLINK (v1.9), incorporating the average dog recombination map (canFam3.1) (*88*) with the --cm-map option. We next ran the adapted snakemake HapNe pipeline for non-humans (https://github.com/currocam/hapne-snakemake) for each population, including all non-related individuals (Figs. 3A) and only those with a >80% ancestry assigned to one genetic cluster in Haplonet analysis (Fig. S20). For each group, we ran it 10 independent times and calculated the median $N_e$ trajectory.

We estimated the exponential growth rate between two time points per population, following the formula: $r = \frac{ln\,(Ne_{t_2})\,-ln(Ne_{t_1})}{t_2 - t_1}$
and expressed as a percentage of change: $(e^r - 1) \cdot 100$ .

We selected three intervals that overlap with known global changes in protection status and their impacts on the demography of European wolves. Assuming a generation time of 4.5 years (*44*) and sampling date of 2020 (Table S2), the first group contained the most recent events (from generation 1 to 10) would overlap the last 45 years (1975-2020); the second group (from generation 10 to 20) would overlap between 90 and 45 years ago (1930-1975); and finally the last group would describe the oldest events (from generation 20 to 50) mostly overlapping the 19$^{th}$ century (1795-1930) (Fig. S19).

**Supplementary Text**

Extended description of Fennoscandian zoo wolves

Clustering analysis showed wolves from Fennoscandian zoos grouping with wild individuals from the Volga-Ural cluster. These wolves came from Ähtäri Zoo, Ahvenniemi field station, Korkeasaari zoo, Ilomantsi, Pohjois-Karjala, Skansen Zoo, and Järvzoo. Their clustering assignment differed from wild individuals in the Fennoscandian region, suggesting that the founders of the zoo populations originally came from Russia.

Limited records are available from the original founders of the populations in those facilities and zoos. Records of wolves acquired by Korkeasaari zoo in Finland (between 1888 and 1918) suggest that they came from the Baltic region or Russia. Some founders from the current wolf population in Skansen Zoo (Sweden), Ähtäri Zoo (Finland) and other Scandinavian zoos might have collected in 1963 from a den in Inari, Lemmenjoki (Finland) (*94*). Additionally, Ähtäri Zoo had 3 breeding female wolves from Russia who were part of the pack. One wolf from Ahvenniemi field station (MW1005) was a wild wolf captured in Ilomantsi (Finland). Since Ilomantsi lies in the easternmost part of Finland, this wolf may have been a long-distance migrant from Russia

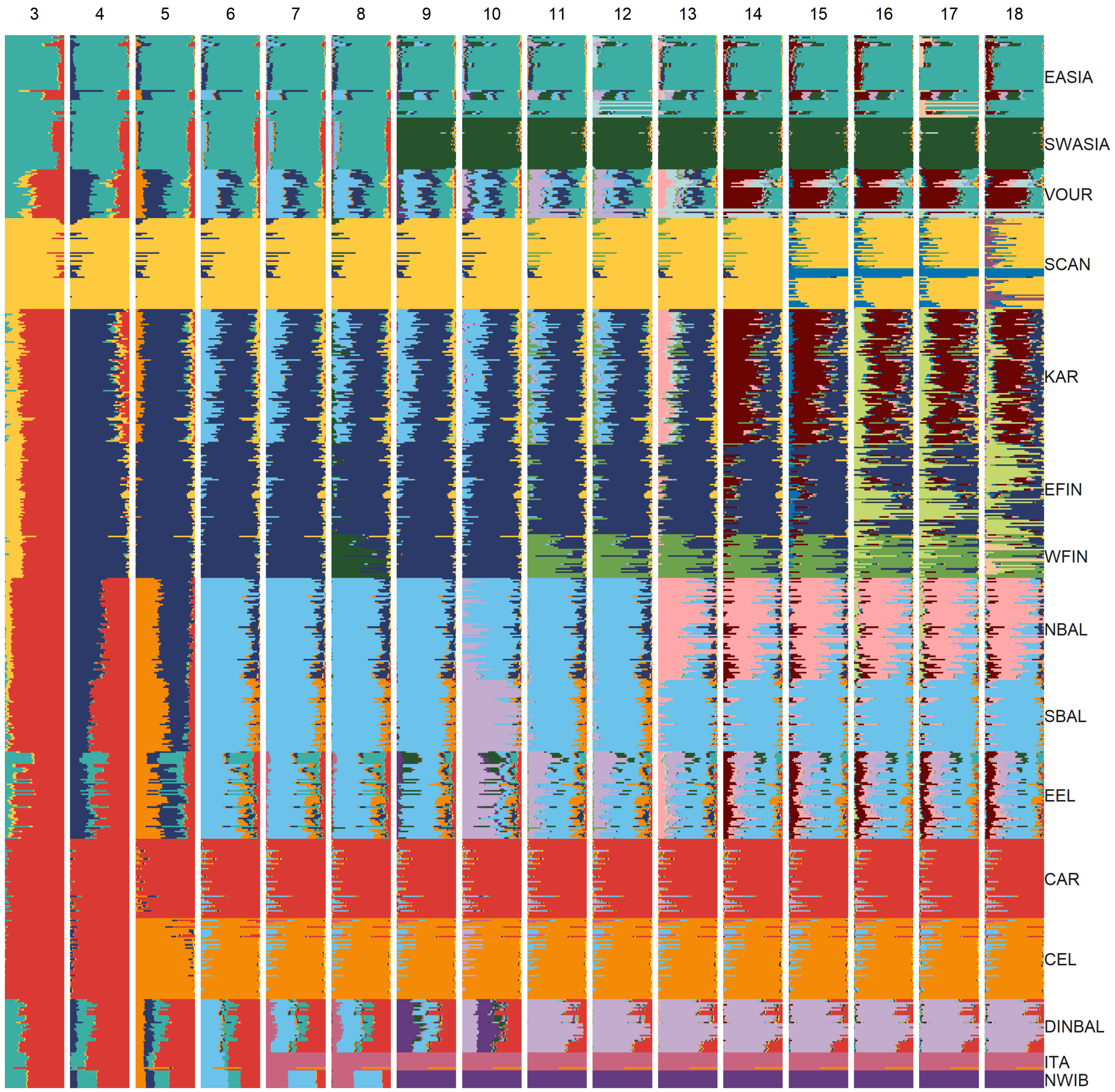


**Fig. S1.** ADMIXTURE (v1.3) (*79*) clustering of unrelated Eurasian wolves (*n*=650) for K values 3-18.

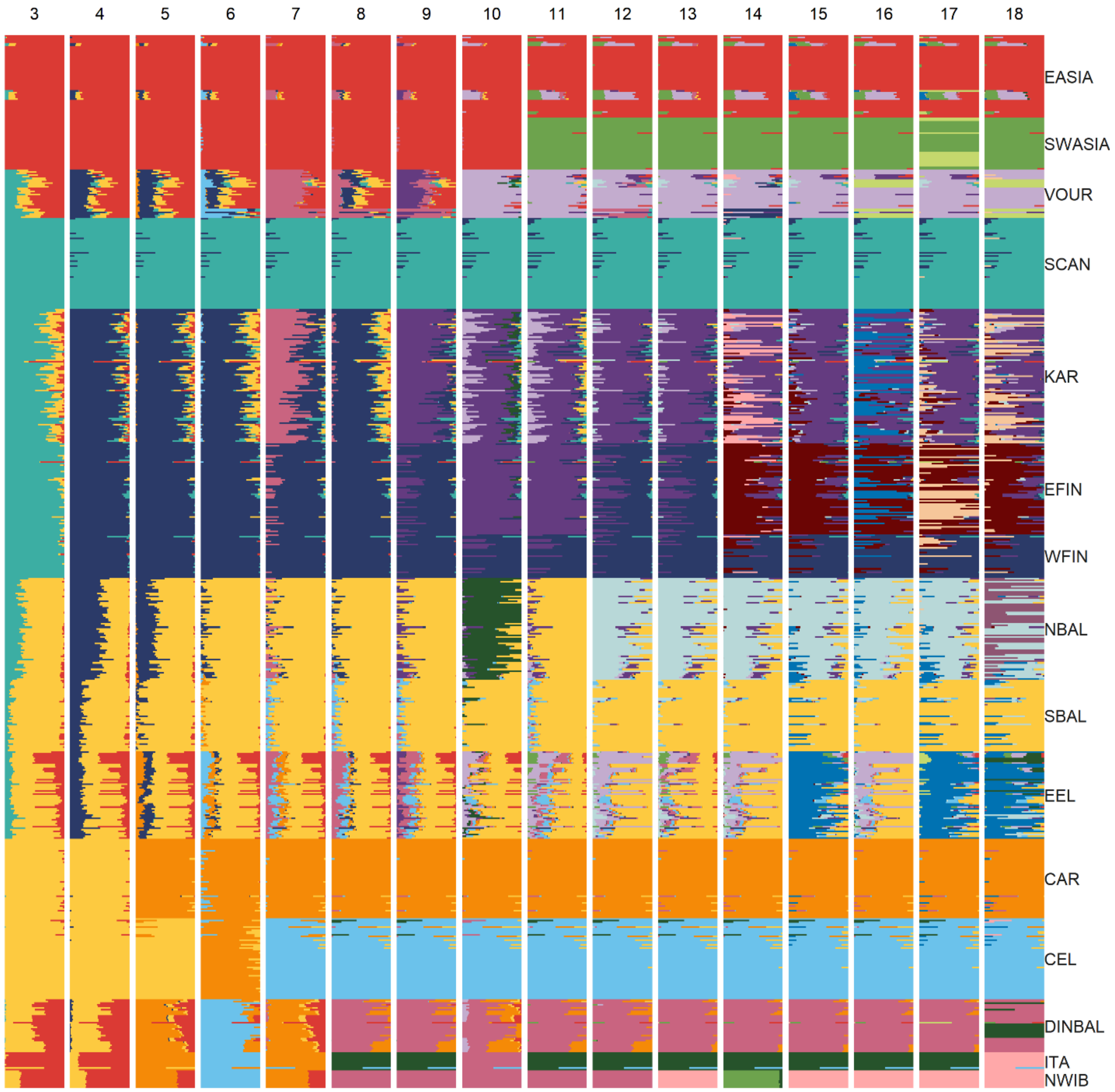


**Fig. S2.** Haplonet admix (v1.0) (*18*) clustering of unrelated Eurasian wolves (*n*=650) for K values 3-18.

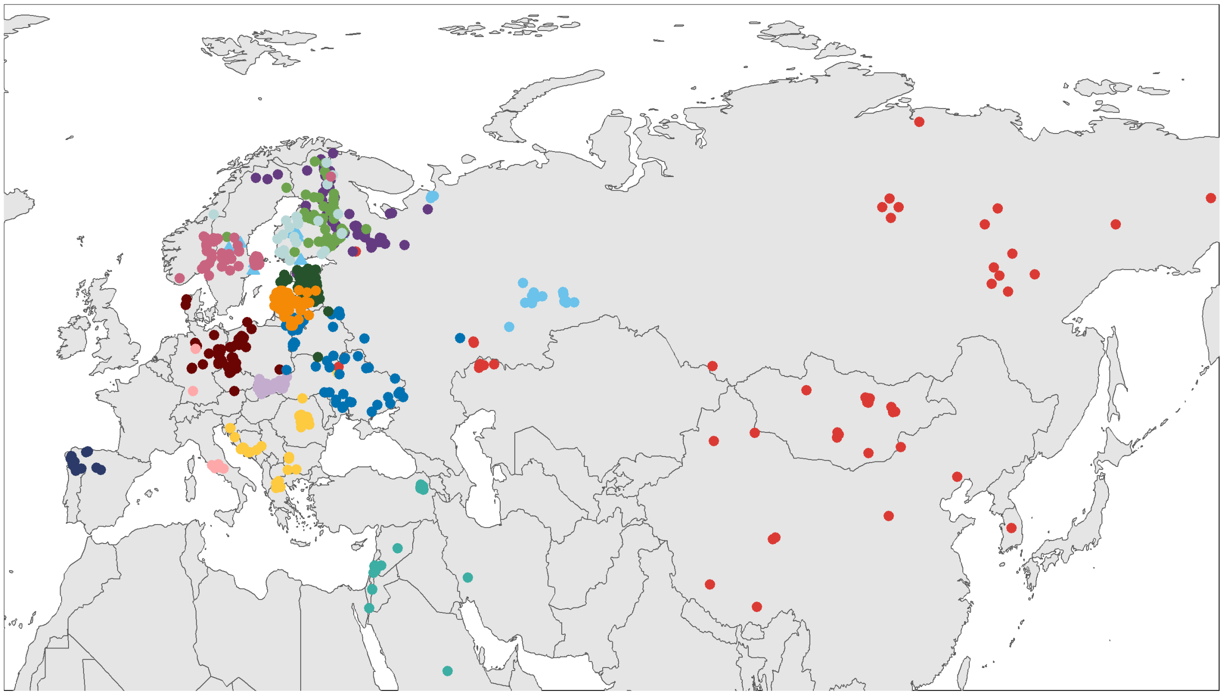

**Fig. S3.** Map of unrelated Eurasian wolves (*n*=650) colored according to the main Haplonet admix cluster assignment (v1, K=15) (*18*).

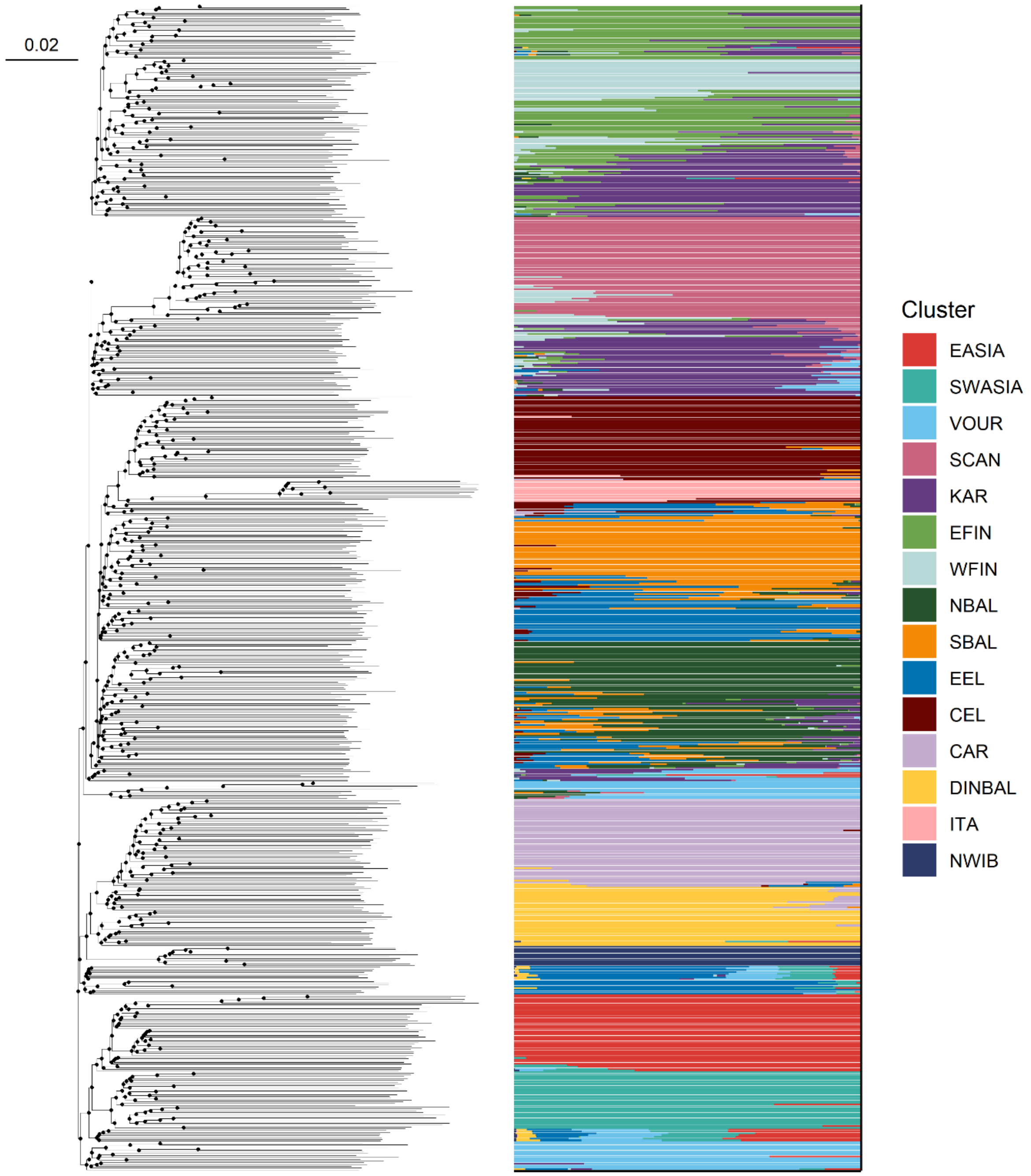


**Fig. S4. Neighbour joining tree and haplotype-based clusters for unrelated Eurasian wolves (*n*=650).** The topology was inferred using Fastme (v2.6.1.3) (*80*) with 100 bootstrap pseudoreplicates. Nodes with more than 90% support are colored black, and the toplogy was midpoint rooted for plotting. The clusters were inferred using Haplonet admix (K=15, v1.0) (*18*).

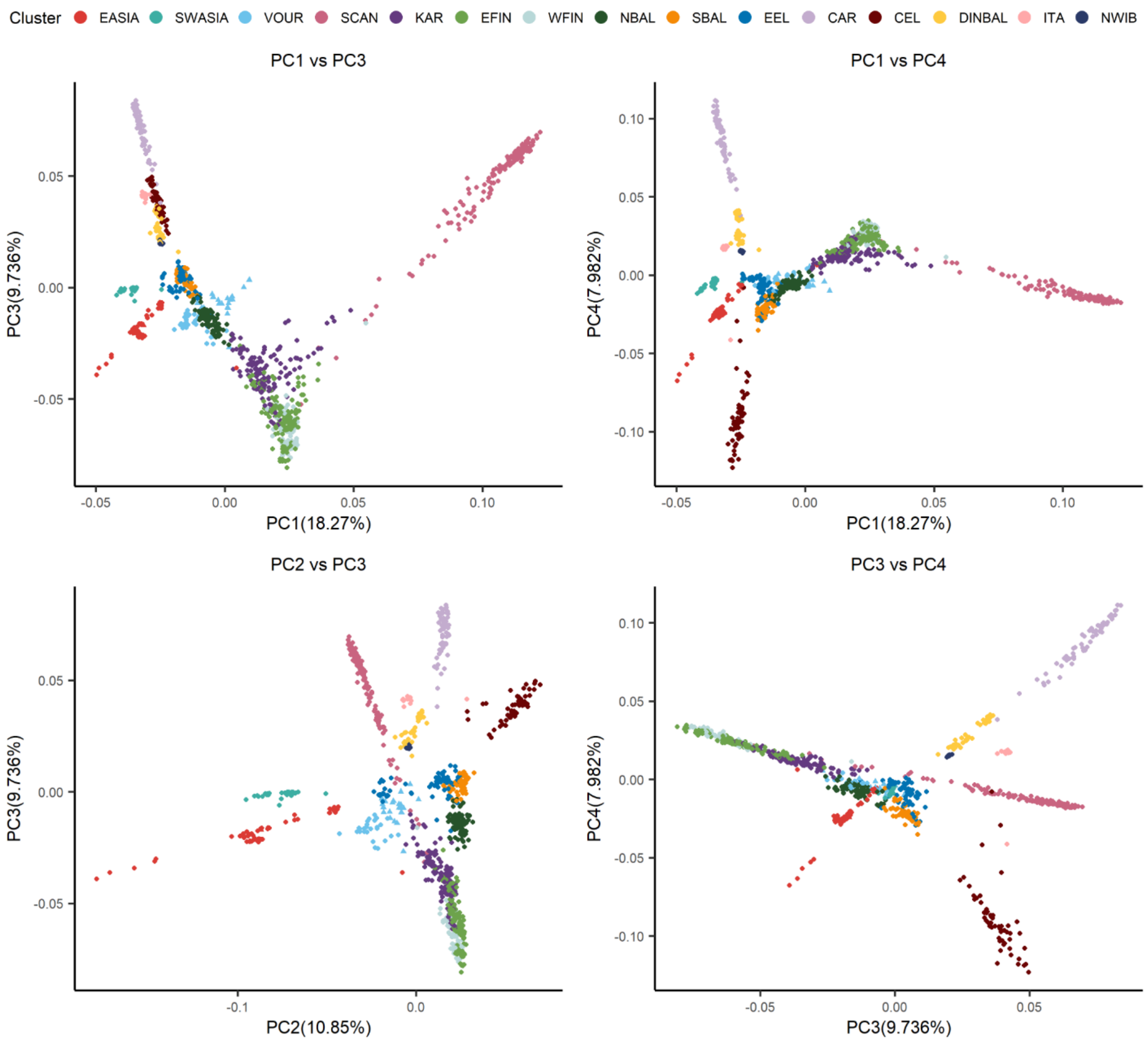


**Fig. S5.** Principal Components Analysis of Eurasian wolves (*n*=1001) using Eigensoft (v8.0) (*19, 81*). The Principal Components (PC's) were estimated using the 650 unrelated wolves, with the related individuals projected. Wolves are colored according to their Haplonet admix cluster (Fig. 1A), and the relationship between PC's 1-4 are shown. The plot of PC1 vs PC2 is found as Fig. 1B.

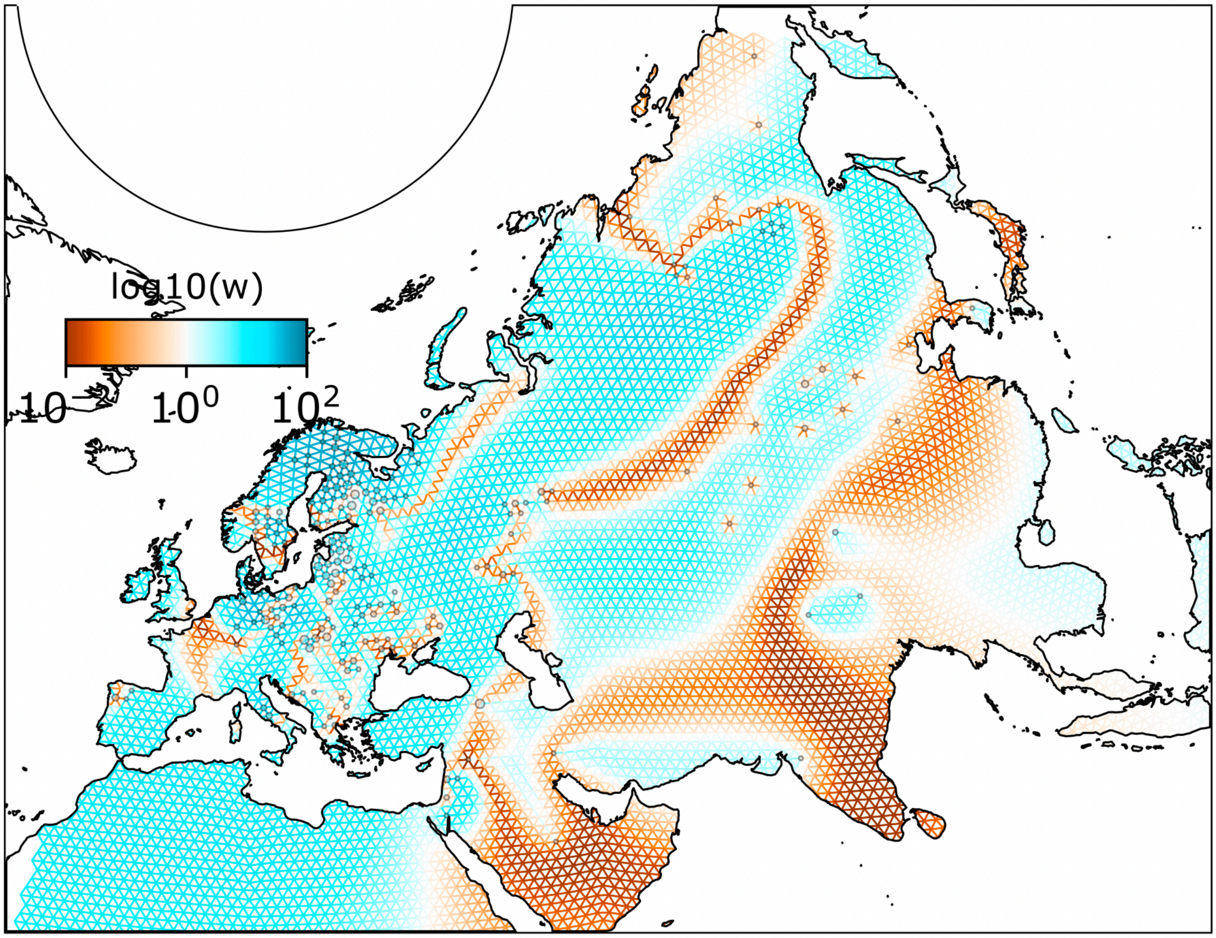


**Fig. S6. Spatial modelling of connectivity amongst unrelated wild Eurasian wolves** (*n*=638) using FEEMS (v1) (*24*). The 12 captive wolves were excluded. Grid corresponds to res = 6 in dgconstruct from the dggridR package, with a cell area of approximately 6,200 km$^2$ and a cell spacing of 110 km. The orange-brown colors represent lower than average effective migration on the log-scale and the blue colors represent higher than average effective migration on the log-scale.

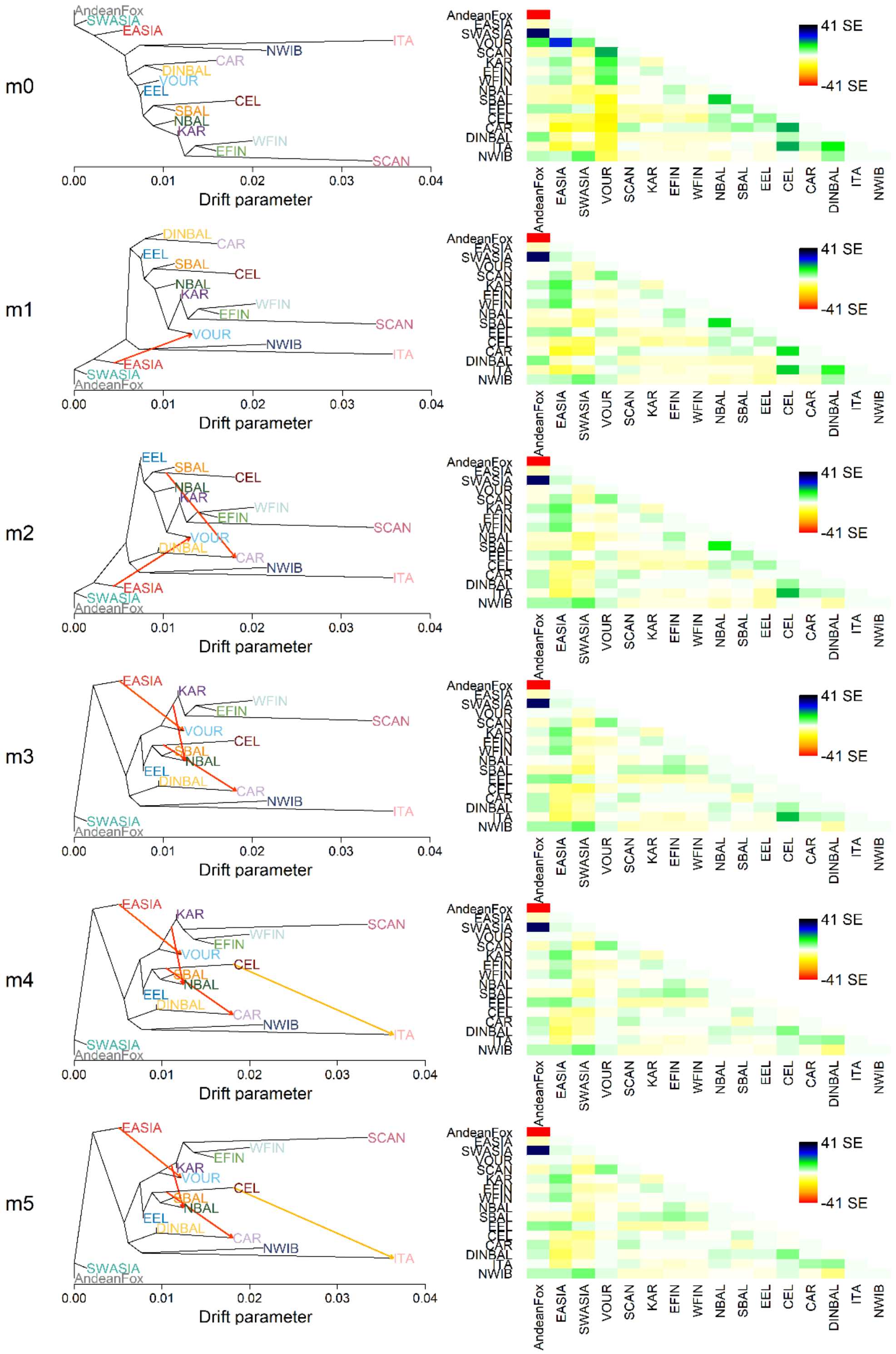


**Fig. S7. Orientagraph (v1.2) (*23*) admixture graph reconstructions for all Eurasian wolf clusters, with 0-5 migration edges (m).** All unrelated wolves from Eurasia (*n*=650) were grouped according to their main cluster assignment using Haplonet admix.

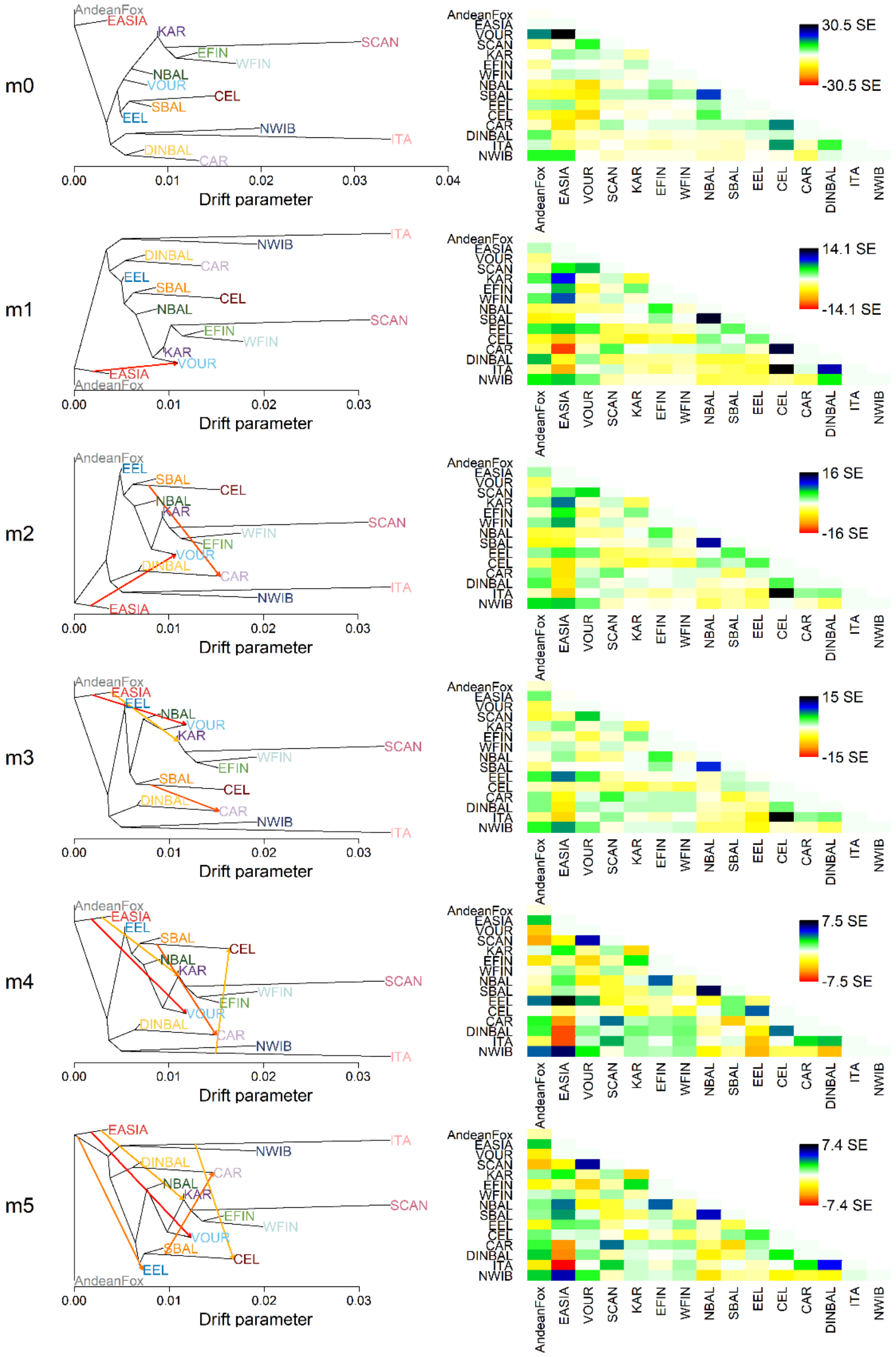


**Fig. S8. Orientagraph (v1.2) (*23*) admixture graph reconstructions for all Eurasian wolf populations, excluding SWASIA, with 0-4 migration edges (m).** All unrelated wolves from Eurasia ($n$=633) were grouped according to their main cluster assignment using Haplonet admix.

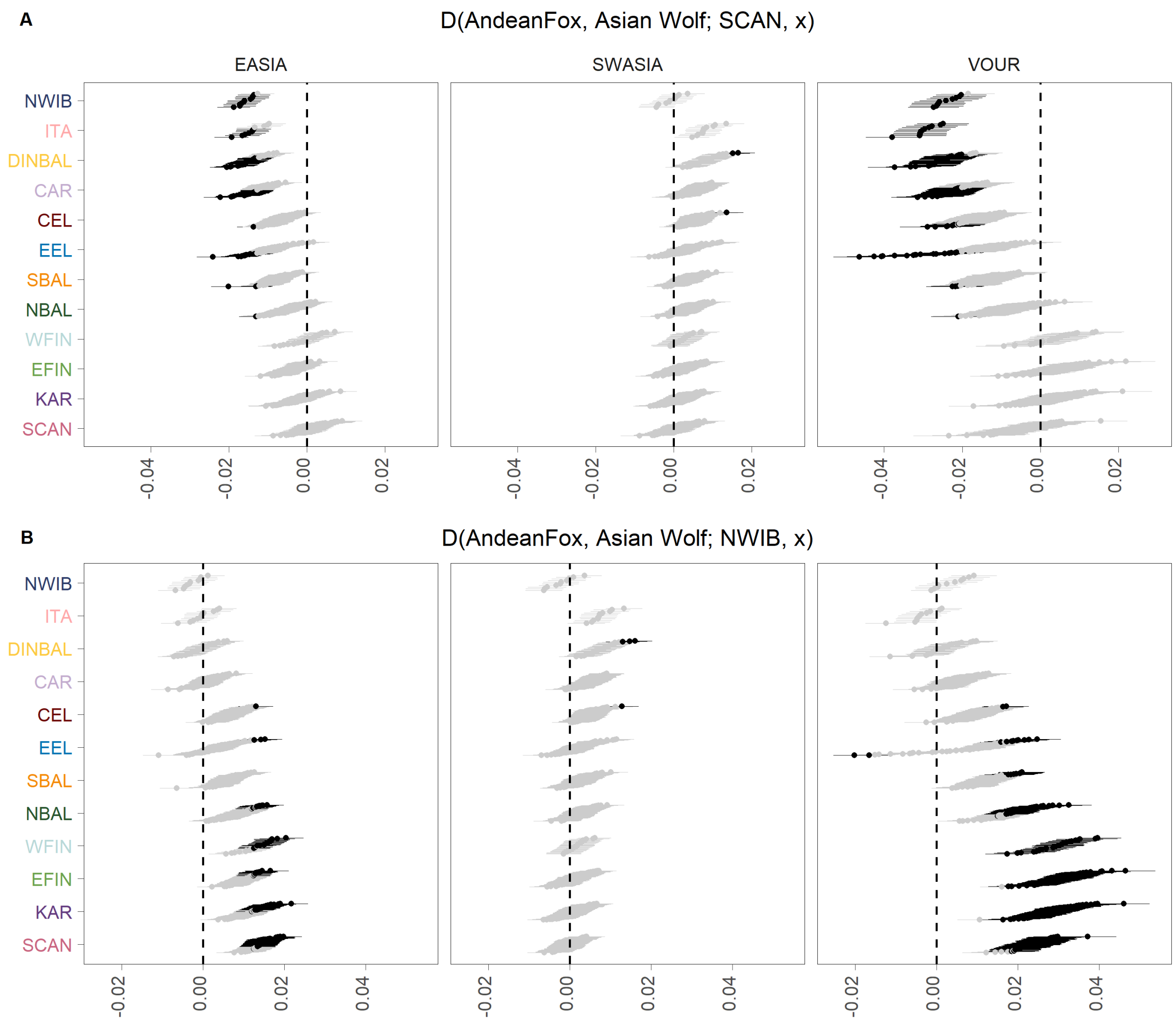


**Fig. S9. D-statistics measuring introgression from Asian wolves into European wolves in the form D(AndeanFox, Asian Wolf; Wolf, x),** estimated using admixtools2 (v2.0.4) (*30*)**,** where x represents each modern European wolf in the panel (*n*=537). The panels on the x axis represents the Asian wolves in the H2 position: EASIA, SWASIA, VOUR. The panels of the y axis place A) a Scandinavian wolf (MW145), and B) a Northwestern Iberian wolf (MW122) in the H3 position. Black points indicate a significant Z-score > |3|.

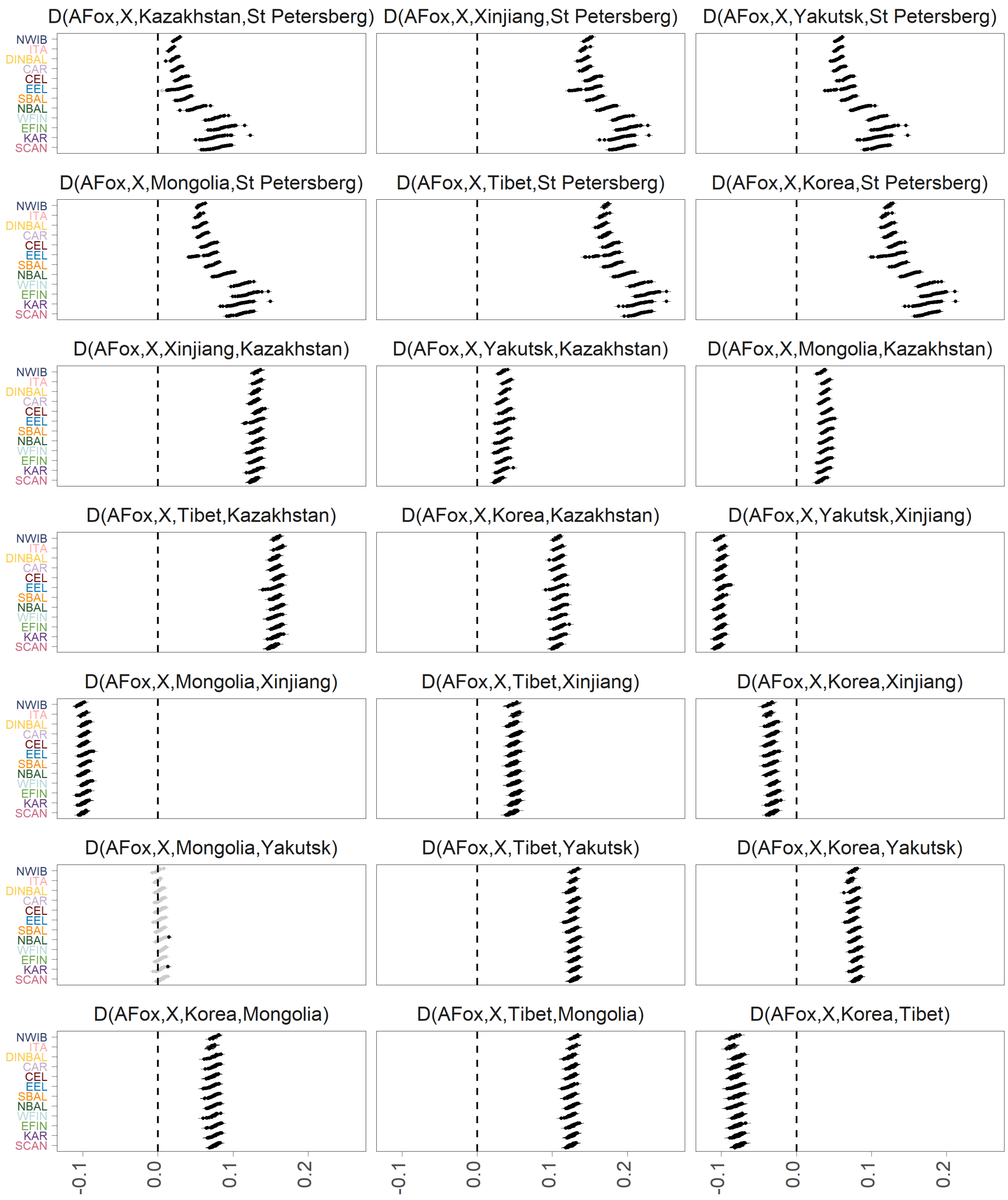


**Fig. S10. D-statistics measuring regional introgression from Asian wolves into European wolves in the form D(AndeanFox, x; Asian Wolf1, Asian Wolf2),** estimated using admixtools2 (v2.0.4) (*30*)**,** where x represents each modern European wolf in the panel (*n*=537). Black points indicate a significant Z-score > |3|.

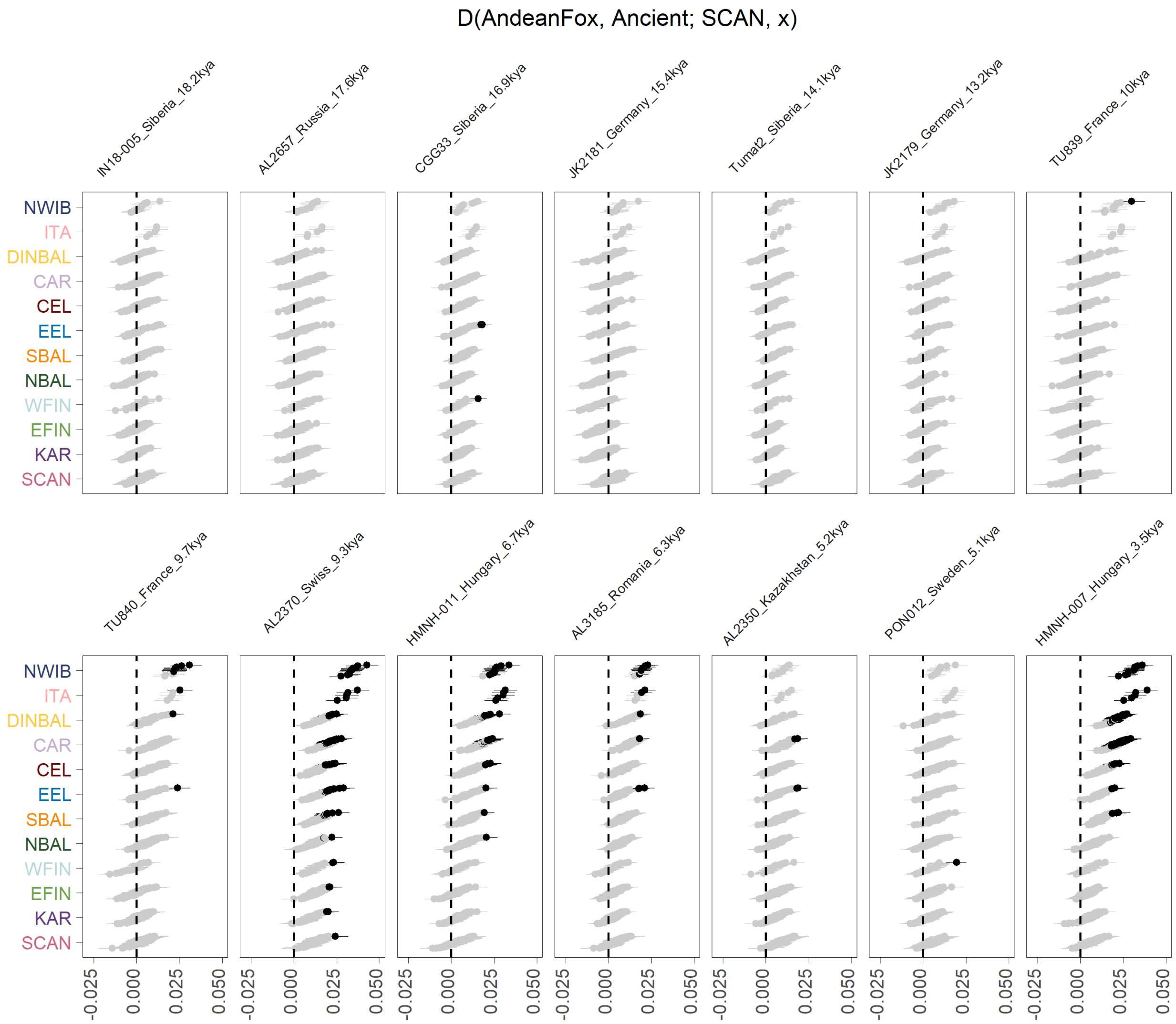


**Fig. S11. D-statistics measuring introgression from ancient wolves into European wolves in the form D(AndeanFox, Ancient; SCAN Wolf, x)**, estimated using admixtools2 (v2.0.4) (*30*), where x represents each modern wolf in the panel (*n*=490). The panels represent the ancient canids in the H2 position. Black points indicate a significant Z-score > |3|.

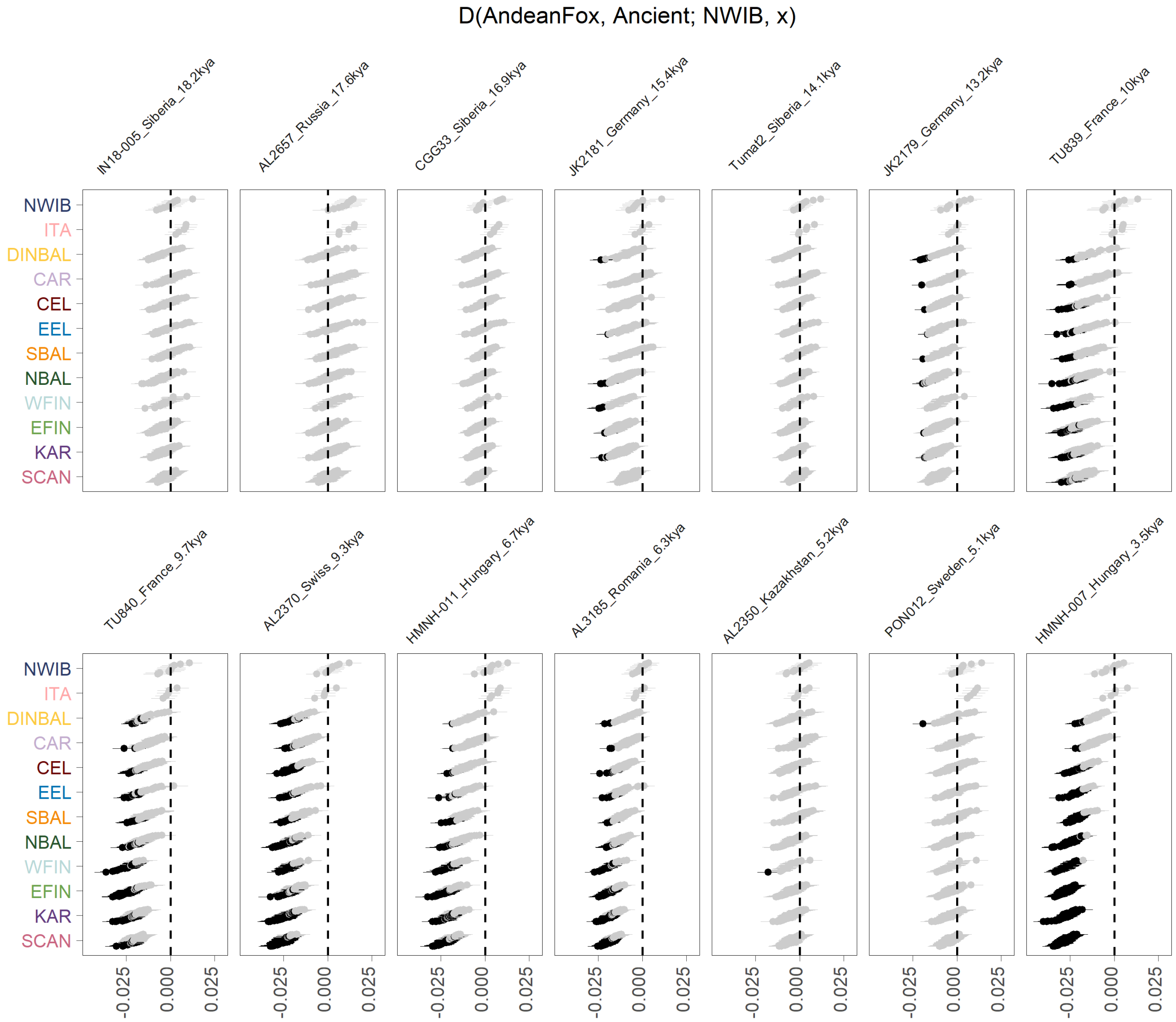


**Fig. S12. D-statistics measuring introgression from ancient wolves into European wolves in the form D(AndeanFox, Ancient; NWIB Wolf, x),** estimated using admixtools2 (v2.0.4) (*30*), where x represents each modern European wolf in the panel (*n*=490), excluding those with significant dog introgression. The panels represent the ancient canids in the H2 position. Black points indicate a significant Z-score > |3|.

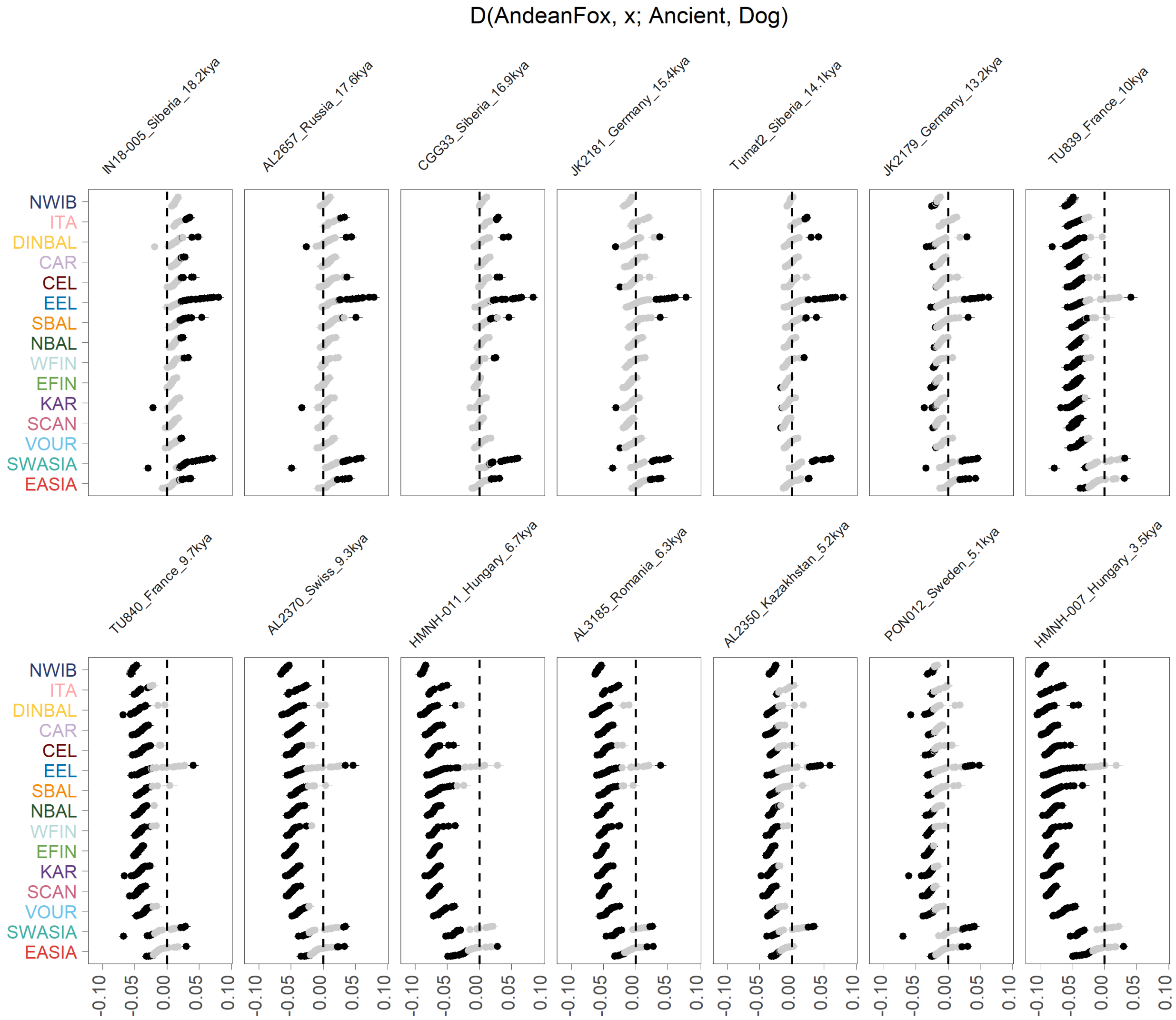


**Fig. S13. D-statistics comparing introgression from dogs to introgression from ancient wolves in the form D(AndeanFox, x; Ancient, Dog),** estimated using admixtools2 (v2.0.4) (*30*), where x represents each modern wolf in the panel (*n*=650). The panels represent the ancient canids in the H3 position, and GShepDog was used for all models in the H4 position. Black points indicate a significant Z-score > |3|.

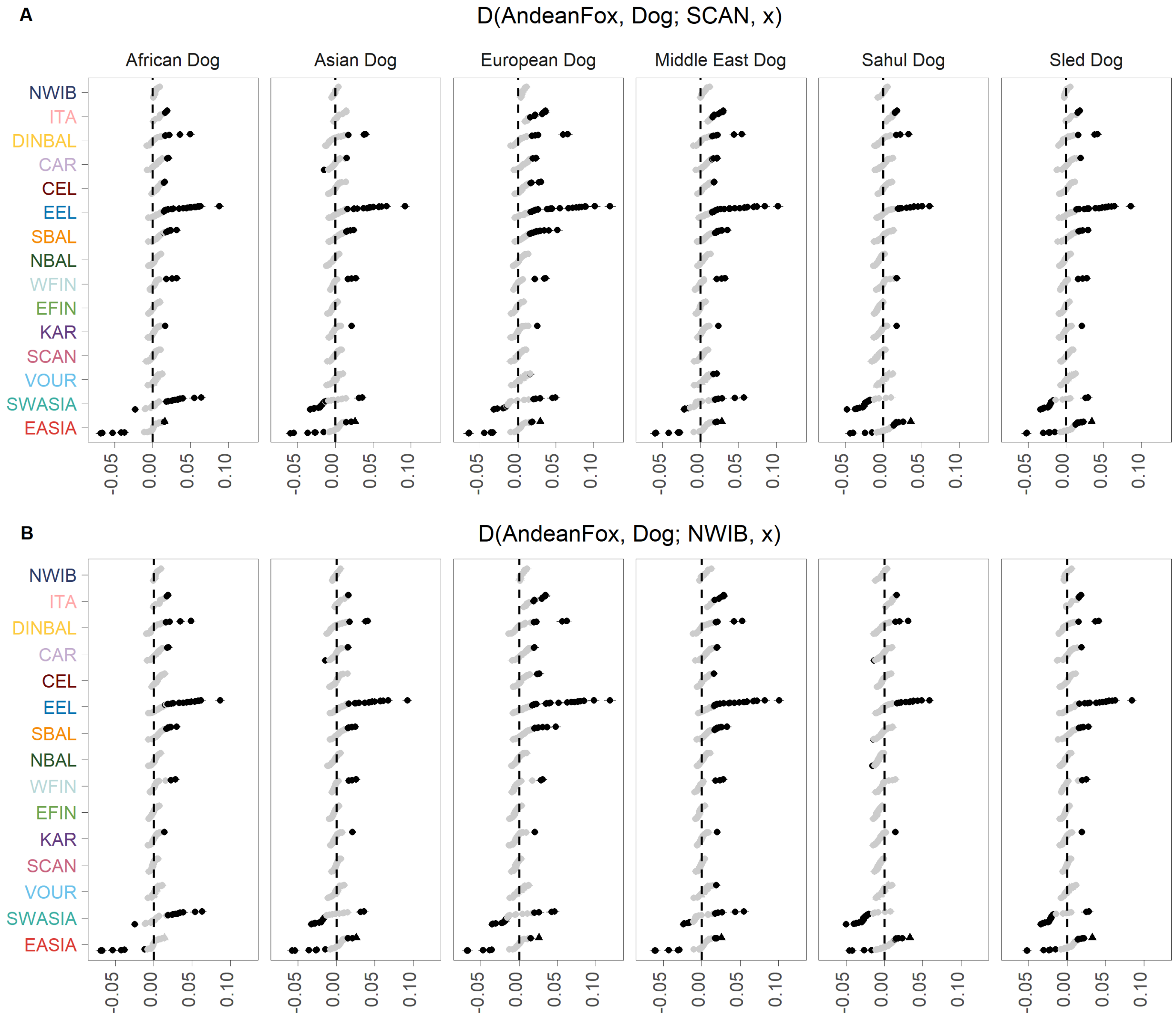


**Fig. S14. D-statistics measuring dog introgression into Eurasian wolves in the form D(AndeanFox, Dog; Wolf, x),** estimated using admixtools2 (v2.0.4) (*30*)**,** where x represents each modern wolf in the panel (*n*=650). The panels on the x axis are the dogs in the H2 position of the D-statistic, representing the major ancestral groups: African Dog, Asian Dog, European Dog, Middle Eastern Dog, Sahul Dog, and Sled Dog. The panels of the y axis place A) a Scandinavian wolf (MW145), and B) a Northwestern Iberian wolf (MW122) in the H3 position. Black points indicate a significant Z-score > |3|.

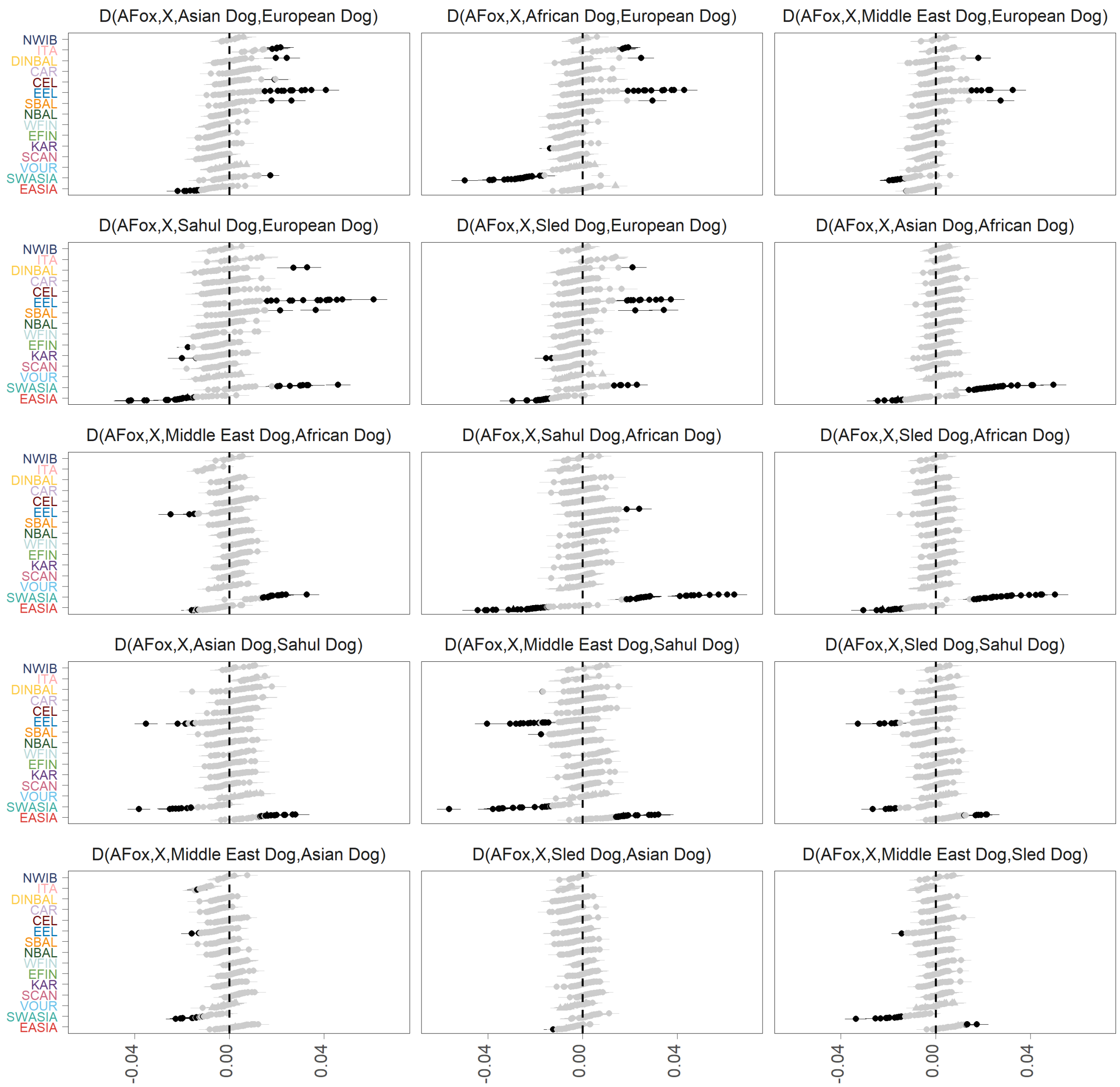


**Fig. S15. D-statistics comparing introgression from dogs of different lineages in the form D(AndeanFox, x; Dog1, Dog2),** estimated using admixtools2 (v2.0.4) (*30*), where x represents each modern wolf in the panel (*n*=650). Black points indicate a significant Z-score > |3|.

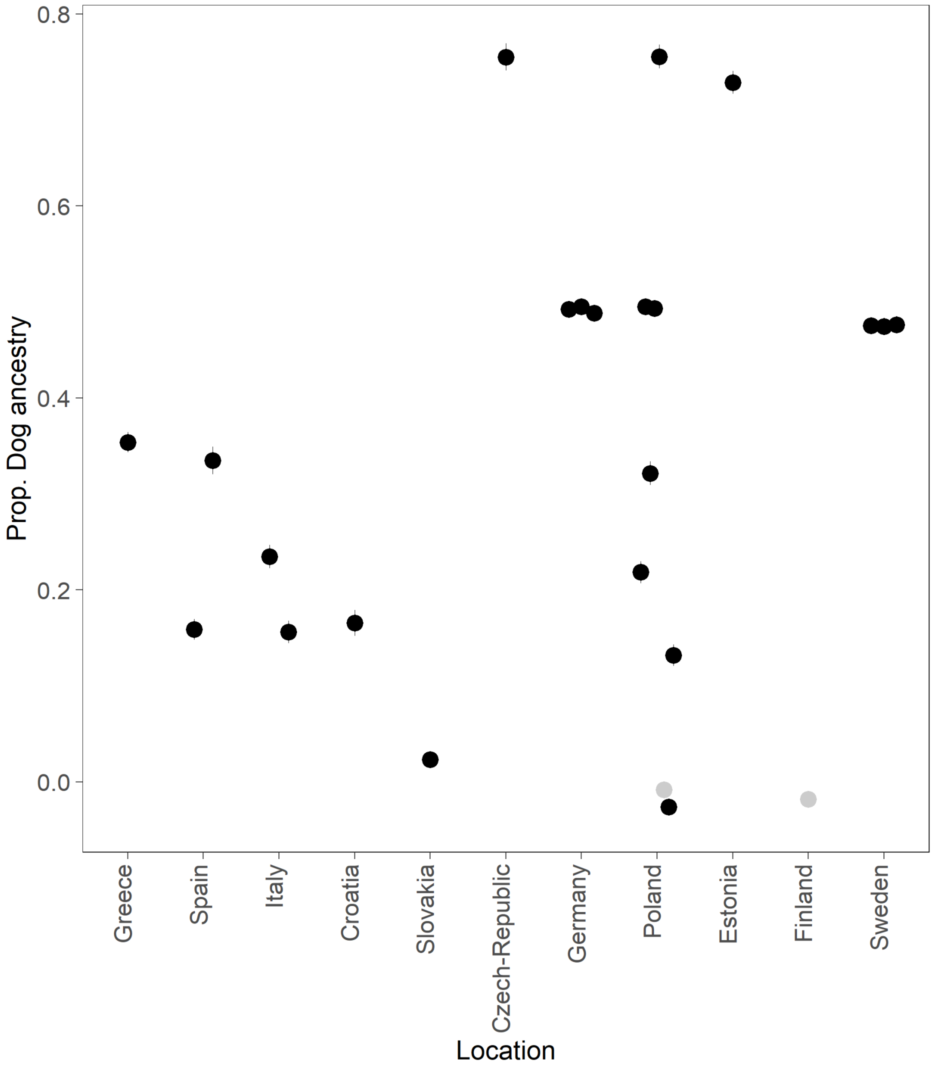


**Fig. S16. $f_4$-ratio tests estimating the proportion of dog ancestry in 24 European hybrids.** $f_4$-ratio tests were calculated in the form *f* 4(FinnishLaplundDog, AndeanFox; x, Wolf)/ *f* 4(FinnishLaplundDog, AndeanFox; GShepDog, Wolf) using admixtools2 (v2.0.4) (*30*). Black points indicate a significant Z-score > |3|. The $f_4$-ratio tests estimating the amount of dog ancestry in European wolves can be found as Fig. 2D.

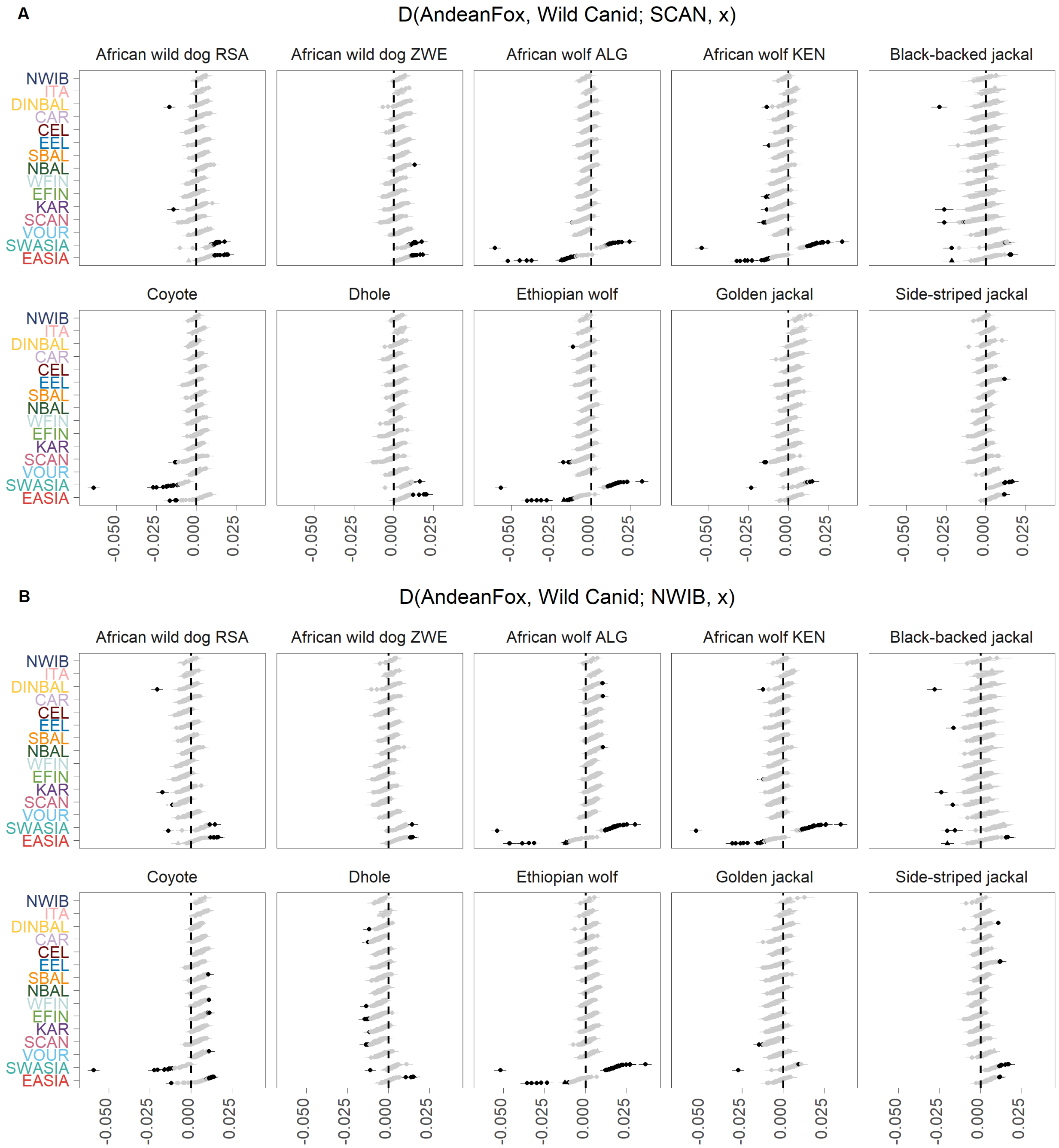


**Fig. S17. D-statistics measuring introgression from wild canids into European wolves in the form D(AndeanFox, Wild; Wolf, x),** estimated using admixtools2 (v2.0.4) (*30*), where x represents each modern wolf in the panel (*n*=650). The panels on the x axis represent the wild canids in the H2 position. The panels of the y axis place A) a Scandinavian wolf (MW145), and B) a Northwestern Iberian wolf (MW122) in the H3 position. Black points indicate a significant Z-score > |3|.

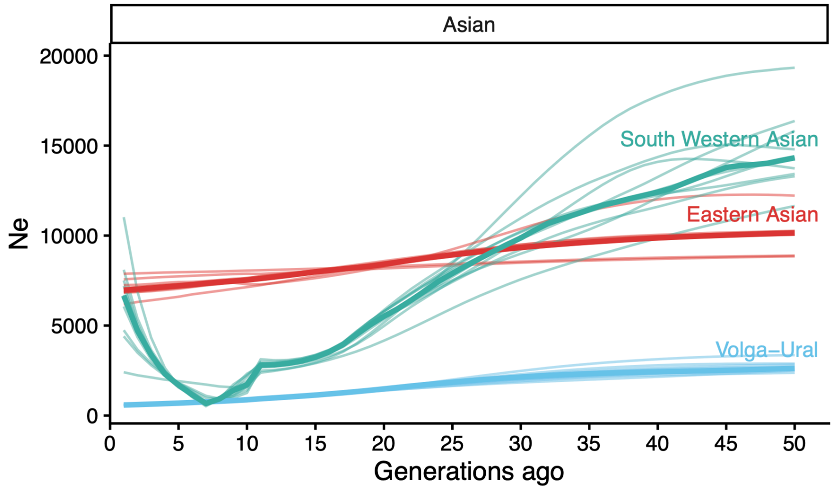


**Fig. S18. Recent demographic history in Asian wolves.** Effective population size trajectories in the last 50 generations (1795-2020), calculated with HapNe-LD (*42*). Thick colored lines represent the median $N_e$ trajectories from 10 independent runs. The light-shaded lines correspond to each independent run.

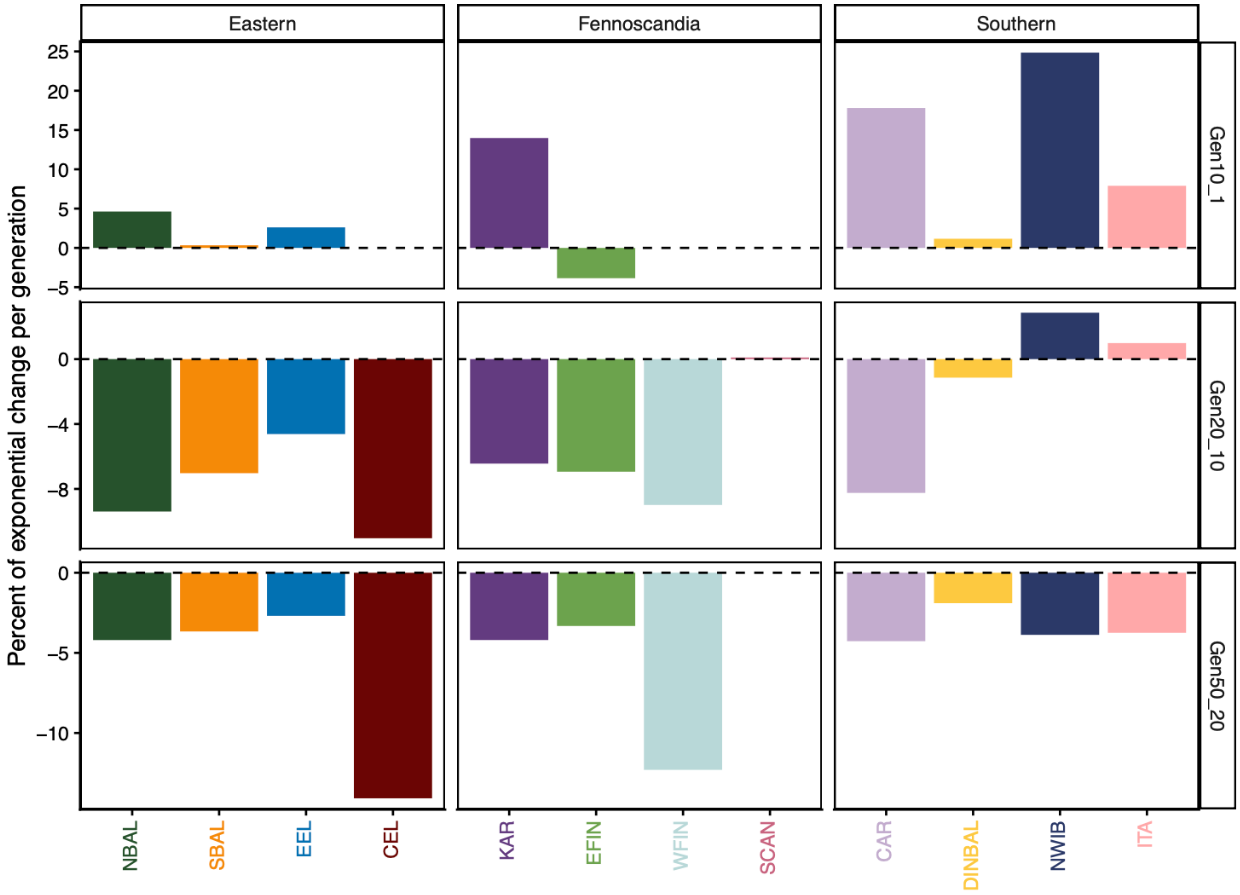


**Fig. S19. Recent demographic $N_e$ trends**, calculated with HapNe-LD as median from 10 independent runs (*42*). The rate of exponential growth for each population is expressed as a percentage. The top panel shows the last 10 generations, which correspond to the last 45 years (1975-2020), assuming a generation time of 4.5 years (*44*). The middle panel (from generation 10 to 20) overlaps between 90 and 45 years ago (1930-1975). The last panel (from generation 20 to 50) overlaps primarily with the 19th century (1795-1930).

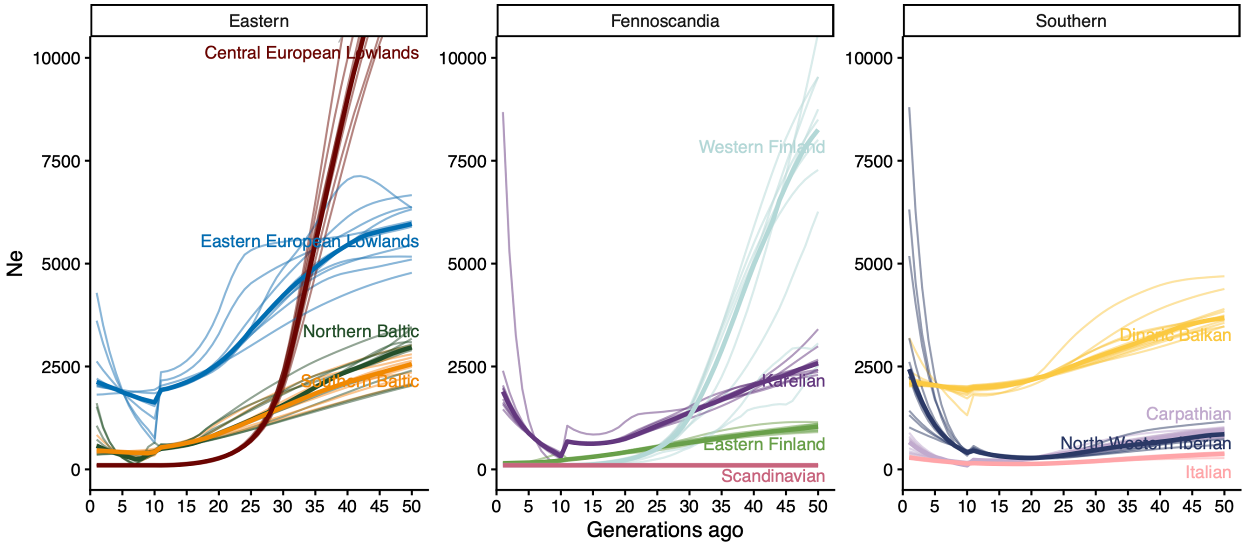


**Fig. S20. Recent demographic $N_e$ trends only including wolves with 80% ancestry assigned to one genetic cluster.** Effective population size trajectories are estimated in the last 50 generations from HapNe-LD (*42*). Thick colored lines represent the median $N_e$ trajectories from 10 independent runs. The light-shaded lines correspond to each independent run.

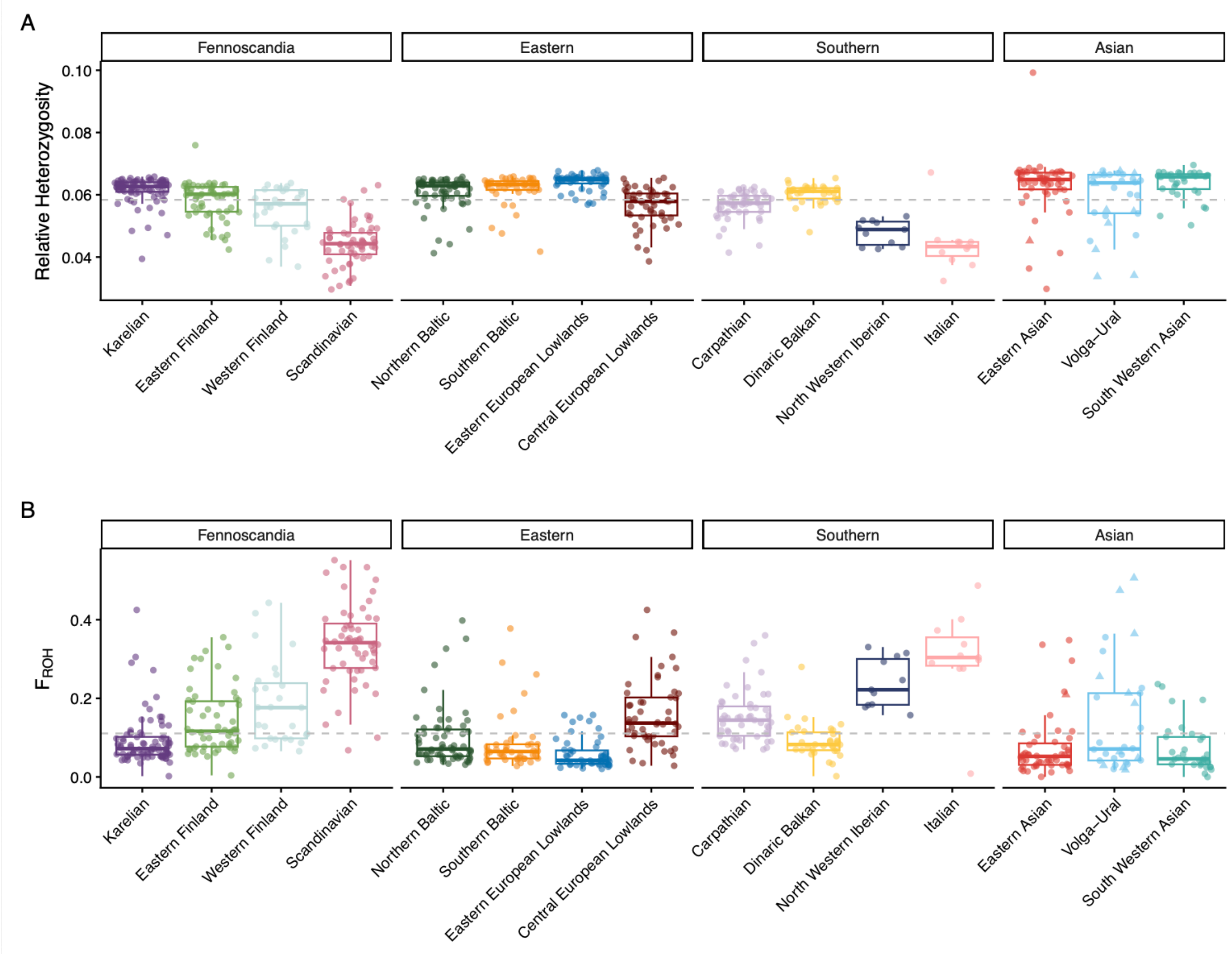


**Fig. S21. Relative heterozygosity and inbreeding in Eurasian wolves.** A) Heterozygosity was estimated as the proportion of heterozygous sites per sample divided by all variants (*n*=52,331,758) (v0.1.17) (*86*). Three samples show excessive heterozygosity compared to other samples from the same group, which show highly admixed profiles on the Haplonet clustering analysis (Fig. S2). The sample MW974, assigned to Eastern Finland, has 51% ancestry from EFIN, 12% from KAR, 19% from EASIA, and 12% from SWASIA. MW097, assigned to the Italian cluster, has 52% ITA and 48% CEL ancestry. MW497, assigned to the Eastern Asian cluster, shows 56% EASIA ancestry, 13% KAR, 20% SWASIA, and 7% VOUR. Dashed grey line represents the average heterozygosity (0.058 het/variants) B) Inbreeding ($F_{ROH}$) was estimated as the proportion of the genome in ROH divided by the autosomal genome using BCFtools roh (v1.20) (*75*). Dashed grey line represents the average $F_{ROH}$=0.11. Triangle shaped points are captive wolves.

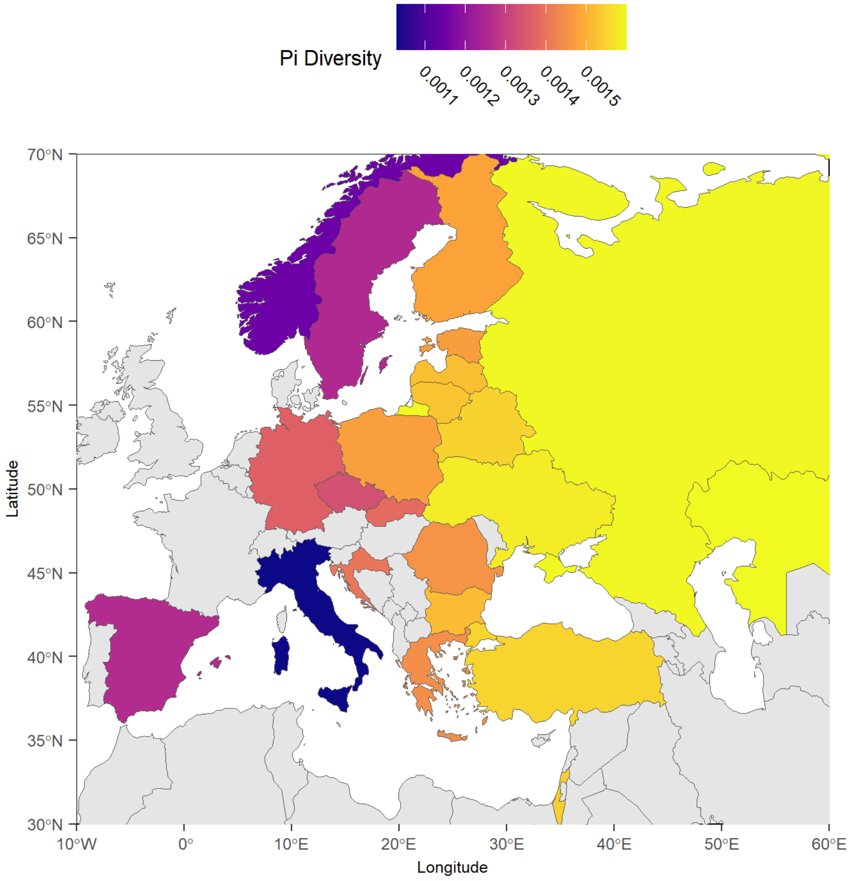


**Fig. S22. Pi-diversity of wolves across Eurasia.** Diversity between wolves was calculated using vcftools (v0.1.17) (*86*) within each country, excluding duplicates. Diversity was not calculated for countries with less than *n*=3.

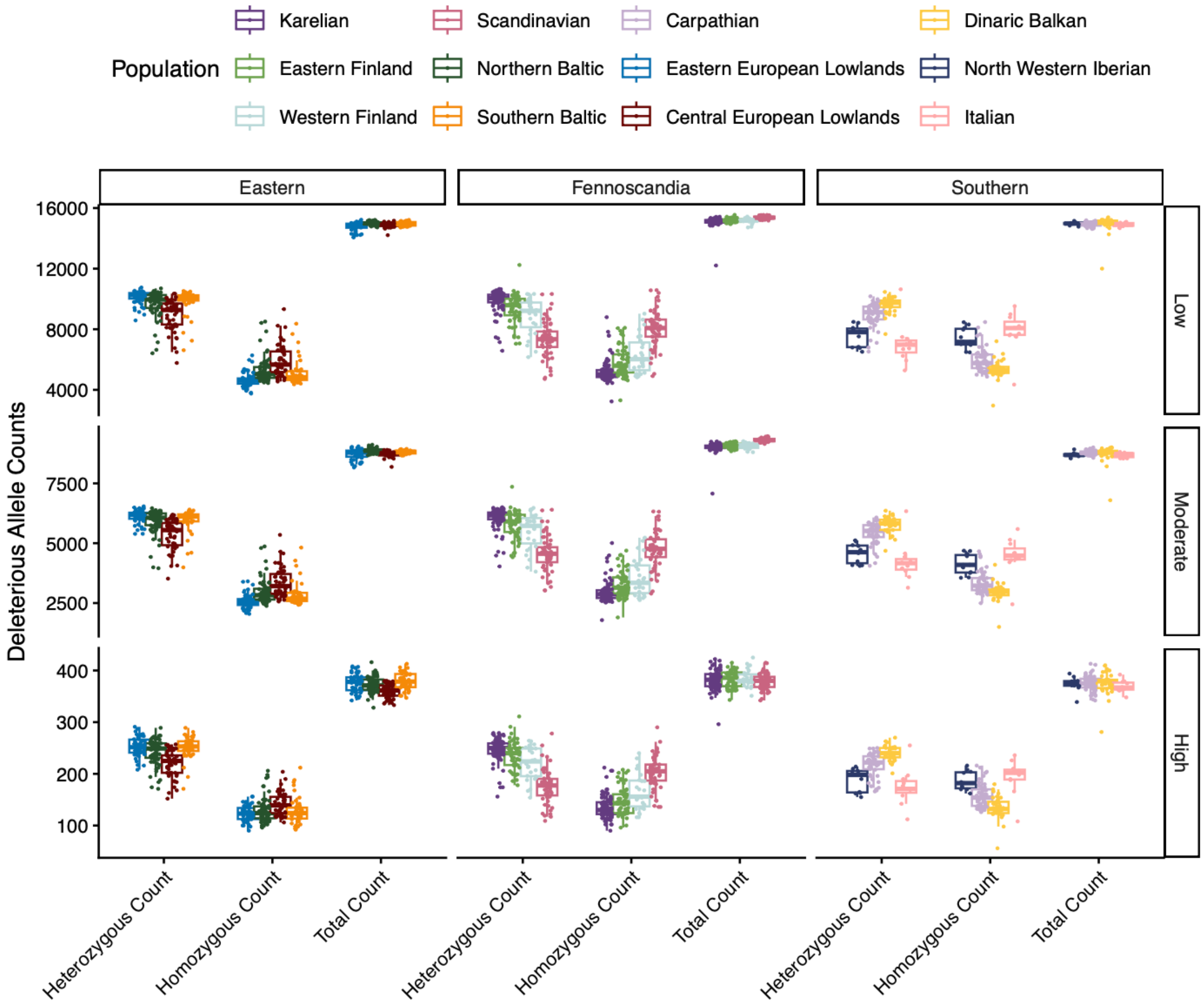


**Fig. S23. Genetic load adding Andean Fox and Dhole also as outgroups for polarization.** Total Deleterious Allele counts (annotated with snpEff (*45*)) for each individual, also segregated by zygosity (homozygous and heterozygous) and deleteriousness (high, moderate, low impact).

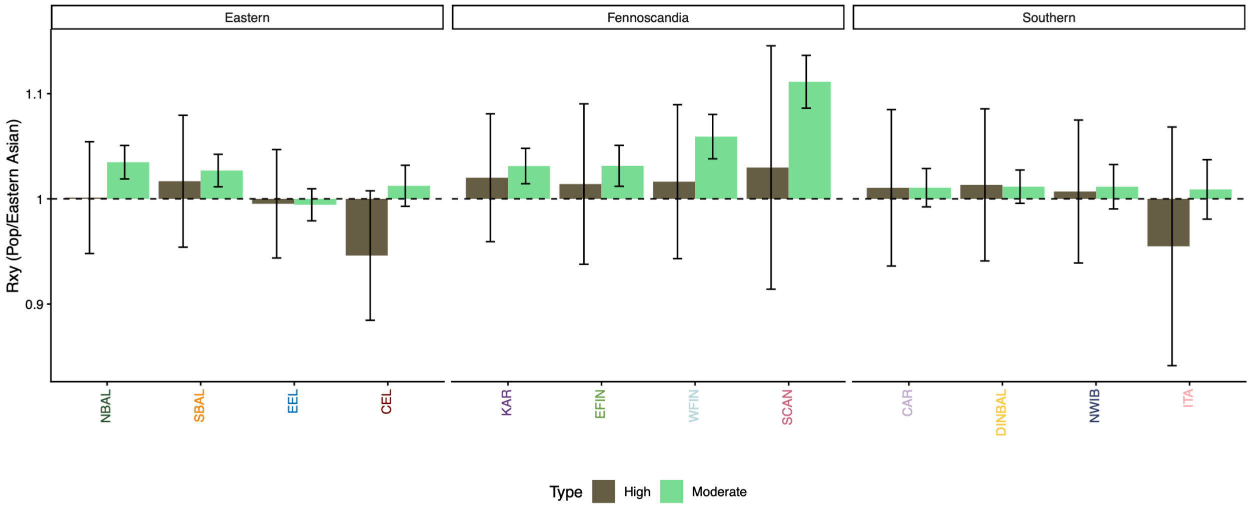


**Fig. S24. Rxy ratio of derived deleterious alleles between each cluster and the Eastern Asian cluster, for high and moderate impact variants (normalized by low Rxy).** Eastern Asian is chosen as Y, because they have the largest and most stable $N_e$ trajectory (Fig. S18). Rxy > 1 indicates a relatively higher frequency of the corresponding category in the studied population, whereas Rxy < 1 indicates a relative frequency deficit. Error bars represent the variance after jackknife resampling of variant sites.

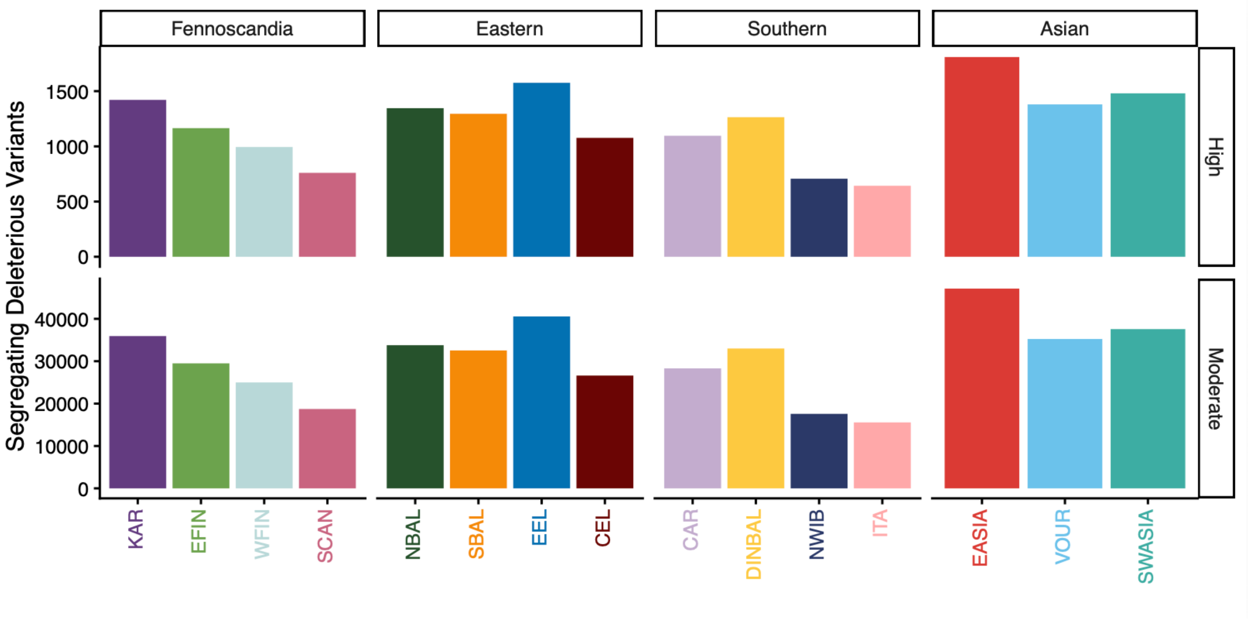


**Fig. S25. Number of segregating deleterious variants per cluster, including Asian wolves.** Segregating variants are those with frequencies between 0 and 1 (not fixed).

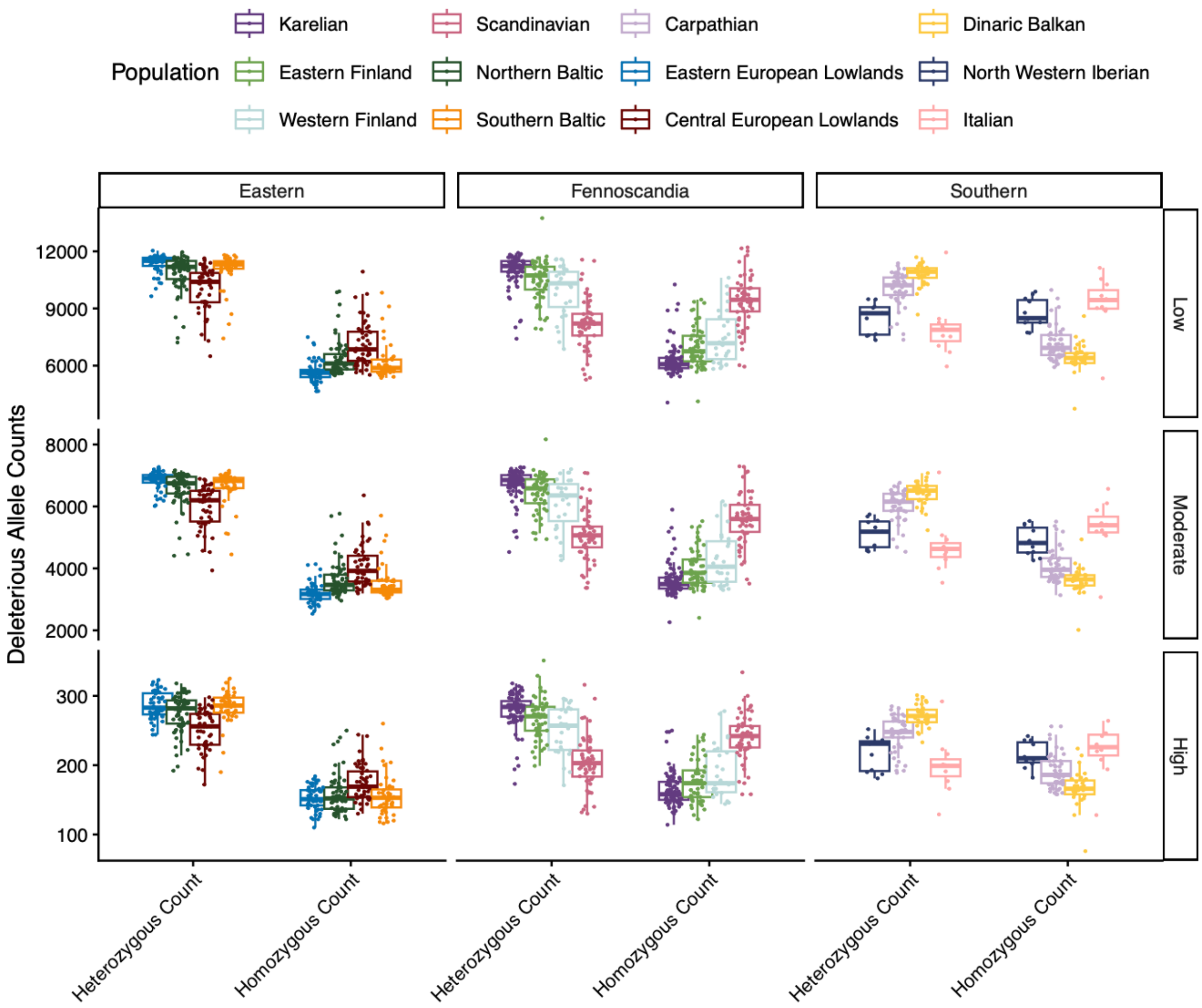


**Fig. S26. Genetic load separated by zygosity state.** Deleterious allele counts (annotated with snpEff) for each individual segregated by deleteriousness (high, moderate, low).

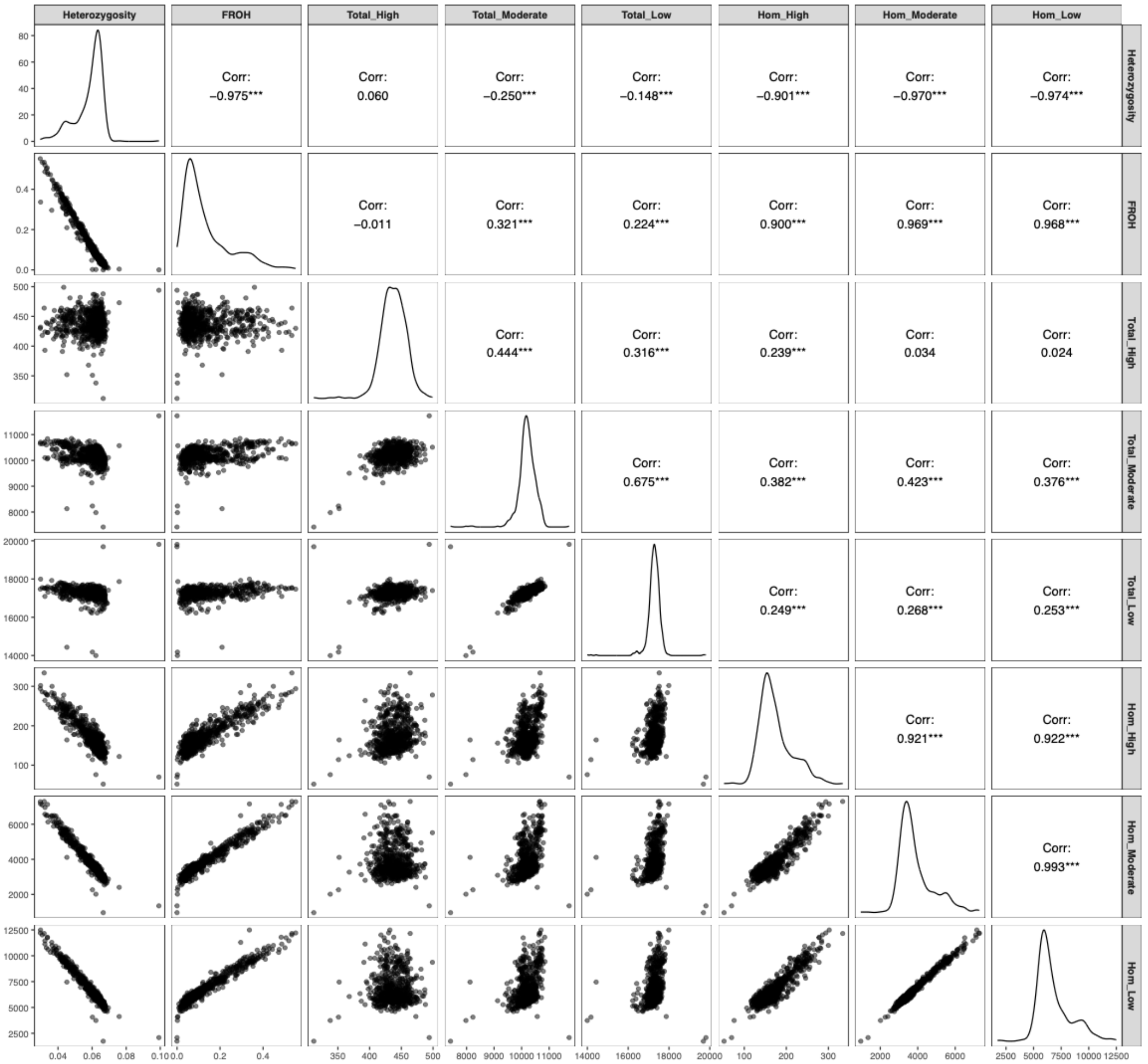


**Fig. S27. Correlations between different summary estimates of diversity, inbreeding and load.** Heterozygosity is estimated as the relative number of heterozygous variants divided by the total genotyped sites. $F_{ROH}$ is the proportion of the genome in ROH. Total_High, Total_Moderate and Total_Low denotes the total allele count in the High, Moderate and Low deleterious category as reported by snpEff. Hom_High, Hom_Moderate and Hom_Low, denote the homozygous allele count in the same categories as for the total count. Stars represent statistical significance: *** p-value<0.001; ** p-value<0.01; and * p-value>0.05.

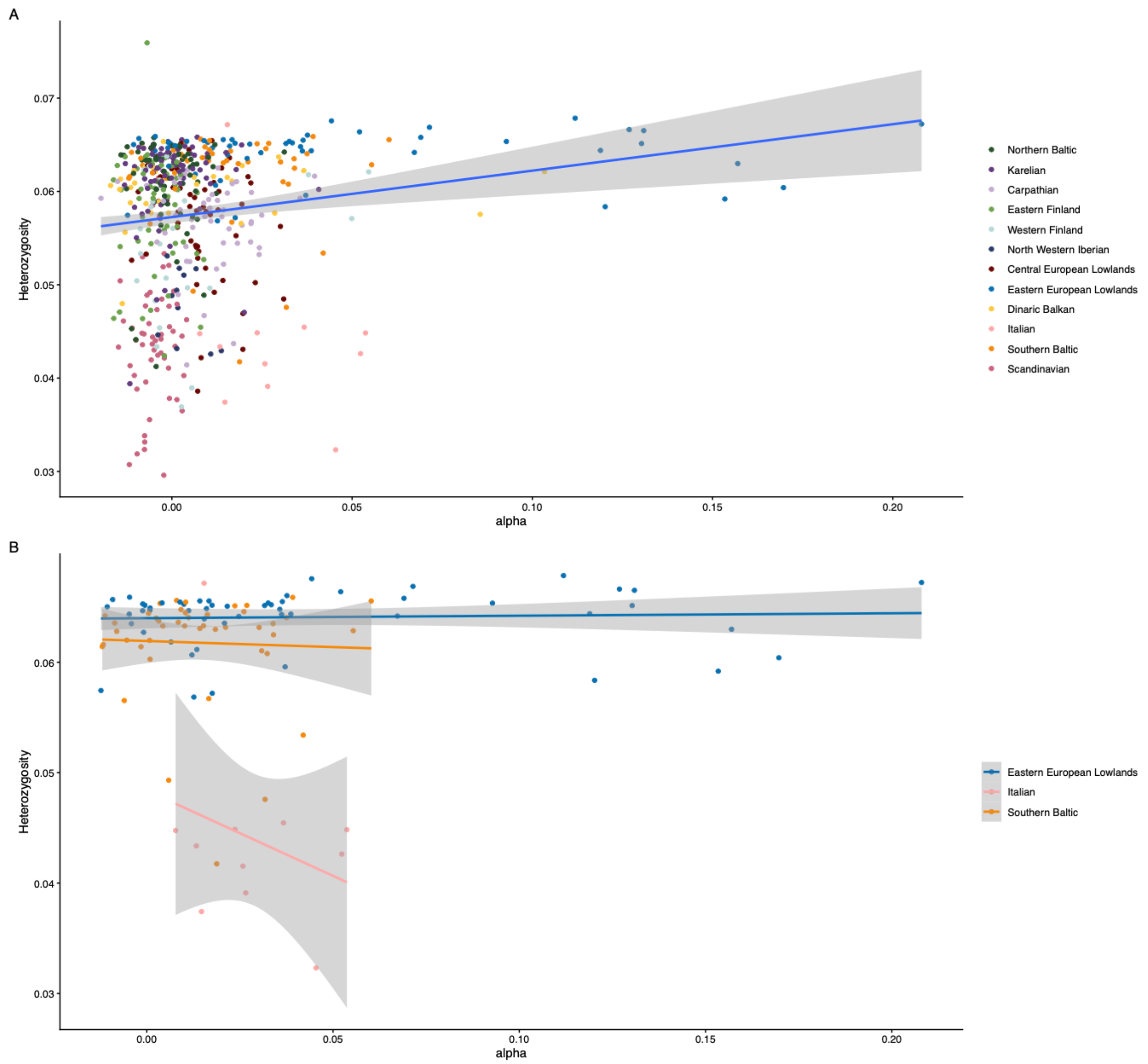


**Fig. S28. Correlation between Heterozygosity and proportion of dog ancestry (alpha).** A) Includes all European wolves. Blue line represents a linear model (Heterozygosity~Alpha) with confidence bands in grey. The linear model accounting for group Heterozygosity~Alpha+Group) has Adjusted R-squared 0.5447 and P-value < 2.2e-16. B) Highlight of the three groups with wolves exhibiting the largest proportion of dog hybridization in their genome. LM (Heterozygosity~Alpha) in Italian group has Adjusted R-squared= -0.0188 and p-value= 0.39; Eastern European Lowlands R-squared= -0.01712 and p-value=0.7437; and Southern Baltic R-squared= -0.02224 and p-value= 0.8009.

**Table S1.** Sample information for all canids imputed in the target panel (*n*=1382). The ID, Museum/Original ID, Biosample Number, Collector/BioProject Number, Collection, Source (public database or generated in this study), Species, Zoo/Wild (whether the sample was from a wild animal or an animal kept in a zoo), Ethical Approval, DNA Quality (modern or degraded), and Location (country) are reported.

**Table S2.** Metadata for Eurasian wolves (*n*=1,068), including Country, Latitude, Longitude and Sample Type (ZWolf = zoo wolves, Wolf = wild wolves, Hybrid = dog-wolf hybrids). Relatedness classification from KING, Haplonet Cluster number, Cluster Name, and Full Cluster Name, Heterozygosity, $F_{ROH}$ (runs of homozygosity, and F4ratio for dog introgression (*f*4(FinnishLaplundDog, AndeanFox; x, Wolf) /*f*4(FinnishLaplundDog, AndeanFox; GShepDog, Wolf)) are reported.

**Table S3.** Sample information for ancient Eurasian wolves screened for use in D-statistics (21 in total, with 14 that had a depth- of coverage of 0.5 used in D-statistics).

| Sample ID | Location | Depth of coverage | Analysis date calBP | Analysis date type | Publication | Used in Dstats |
|---|---|---|---|---|---|---|
| CGG19 | Mus-Khaya, Siberia | 0.089 | 19710 | Radiocarbon | (*61*) | no |
| IN18-005 | Belaya Gora, Siberia | 1.867 | 18148 | Radiocarbon | (*61*) | yes |
| AL2657 | Eliseevichi, western Russia | 0.947 | 17633 | Radiocarbon | (*61*) | yes |
| CGG33 | Ulakhan Sular, Siberia | 16.464 | 16863.5 | Radiocarbon | (*32*) | yes |
| JK2181 | Hohle Fels, Germany | 0.587 | 15447 | Radiocarbon | (*61*) | yes |
| Tumat1 | Tumat, Siberia | 0.472 | 14618 | Radiocarbon | (*61*) | no |
| Tumat2 | Tumat, Siberia | 5.943 | 14122 | Radiocarbon | (*17, 23*) | yes |
| CGG21 | Berelekh, Siberia | 0.221 | 13974.5 | Radiocarbon | (*61*) | no |
| CGG20 | Nikita Lake site, Muksunuokha River, Siberiariver | 0.089 | 13660 | Radiocarbon | (*61*) | no |
| 367 | Ballynamintra Cave, Ireland | 0.008 | 13334 | Radiocarbon | (*61*) | no |
| JK2179 | Hohle Fels, Germany | 2.712 | 13229 | Radiocarbon | (*61*) | yes |
| IRK | Plunkett Cave, Kesh caves, Ireland | 0.234 | 13219 | Radiocarbon | (*61*) | no |
| TU839 | Le Parc du Château, Auneau, France | 0.529 | 10055 | Radiocarbon | (*61*) | yes |
| TU840 | Le Parc du Château, Auneau, France | 1.074 | 9715.5 | Radiocarbon | (*61*) | yes |
| AL2370 | Dürriloch, Boltigen i.S., Bern, Switzerland | 0.783 | 9348 | Radiocarbon | (*61*) | yes |
| CH1109 | Noyen-Sur-Seine, France | 0.029 | 8629 | Radiocarbon | (*61*) | no |
| DS-CANIS-HMNH-011 | Leány-barlang, Hungary | 1.619 | 6719 | mtDNA tip dating | (*61*) | yes |
| AL3185 | Pietrele, Romania | 4.252 | 6307 | Radiocarbon | (*61*) | yes |
| AL2350 | Botai, Kazakhstan | 2.009 | 5169 | Radiocarbon | (*61*) | yes |
| PON012 | Alvastra, Sweden | 2.317 | 5079 | Contextual | (*61*) | yes |
| CANIS-HMNH-007 | Istállóskő Cave, Hungary | 1.234 | 3525 | mtDNA tip dating | (*61*) | yes |

**Table S4. Proportion of dog genetic material in European wolves (*n*=873),** measured using *f* 4-ratio statistics. Average dog ancestry is calculated with non-significant values set to 0.

| Cluster | Proportion of individuals with dog ancestry | Average proportion of dog ancestry in the genome | *n* |
|---|---|---|---|
| SCAN | 0.0073 | 0.0003 | 137 |
| KAR | 0.0073 | 0.0003 | 137 |
| EFIN | 0 | 0 | 146 |
| WFIN | 0.0984 | 0.0044 | 61 |
| NBAL | 0.0127 | 0.0004 | 79 |
| SBAL | 0.2909 | 0.0108 | 55 |
| EEL | 0.4559 | 0.0369 | 68 |
| CEL | 0.0545 | 0.0016 | 55 |
| CAR | 0.0806 | 0.0027 | 62 |
| DINBAL | 0.1429 | 0.0081 | 35 |
| ITA | 0.4667 | 0.0210 | 15 |
| NWIB | 0 | 0 | 10 |

**Table S5. Effective population sizes ($N_e$) and trends in Eurasian wolves.** Estimates are derived from HapNe-LD, from the median of 10 independent runs. CurrentNe represents $N_e$ at timepoint 1 and minNe and maxNe represent the minimum and maximum $N_e$ reach through the last 50 generations, respectivley. The percentage of exponential rate changes is obtained per generation (see Methods) and is shown at different time bins: Gen50_20 (1795-1930), Gen20_10 (1930-1975), Gen_10_1 (1975-2020) and Gen_50_10 (1795-1975).

| | | | | Percentage exponential rate change per generation | | | |
|---|---|---|---|---|---|---|---|
| Cluster | minNe | maxNe | Current Ne | Gen_50_20 | Gen_20_10 | Gen_10_1 | Gen_50_10 |
| Karelian | 390 | 2980,5 | 1513 | 4.20 | 6.44 | -13.98 | 1.26 |
| Eastern Finland | 154 | 1288 | 154 | 3.32 | 6.94 | 3.86 | 5.11 |
| Western Finland | 99,5 | 8535 | 100 | 12.30 | 8.99 | 0 | 11.46 |
| Scandinavian | 99 | 100 | 100 | 0 | -0.10 | 0 | -0.03 |
| Northern Baltic | 250,5 | 3173 | 554,5 | 4.20 | 9.39 | -4.63 | 4.36 |
| Southern Baltic | 405,5 | 3078 | 453 | 3.67 | 7.03 | -0.34 | 4.42 |
| Carpathian | 117,5 | 1154,5 | 686 | 4.27 | 8.25 | -17.80 | 0.71 |
| Eastern European Lowlands | 1631 | 6187 | 2070 | 2.69 | 4.62 | -2.61 | 2.56 |
| Central European Lowlands | 99 | 17938 | 100 | 14.08 | 11.04 | 0 | 13.31 |
| Dinaric Balkan | 1922 | 4242 | 2135,5 | 1.90 | 1.14 | -1.16 | 1.44 |
| Northwestern Iberian | 282 | 4931 | 4931 | 3.88 | -2.86 | -24.85 | -4.21 |
| Italian | 125,5 | 441,5 | 300 | 3.75 | -0.99 | -7.90 | 0.66 |
| Eastern Asian | 6963,5 | 10205 | 6963,5 | 0.63 | 1,11 | 0,87 | 0.94 |
| Volga-Ural | 587,5 | 2860,5 | 587,5 | 1.93 | 5.39 | 4,55 | 3.81 |
| Southwestern Asian | 670,5 | 15570 | 6666,5 | 3.23 | 12.39 | -14.00 | 1.93 |
| Average (Exclude Italian and Northwestern Iberian) | | | | 5,06 | 6.37 | 3.67 | 4.46 |
| Average | | | | 4.85 | 4.99 | -5.79 | 3.42 |

**Table S6. Average genetic load (allele counts) for moderate- and high-impact variants per population.** Ratio and percentage change relate each cluster to the average of all European wolves.

| | Moderate Impact Variants | | | High Impact Variants | | |
|---|---|---|---|---|---|---|
| Group | Ratio from average | Percent change from average | Average Count Deleterious Alleles | Ratio from average | Percent change from average | Average Count Deleterious Alleles |
| Karelian | 1.01 | 0.91 | 10344.2 | 1.01 | 1.29 | 443.8 |
| Eastern Finland | 1.02 | 1.61 | 10415.9 | 1.01 | 1.27 | 443.8 |
| Western Finland | 1.02 | 1.54 | 10408.7 | 1.00 | 0.42 | 440.0 |
| Scandinavian | 1.04 | 3.97 | 10658.0 | 1.01 | 1.19 | 443.4 |
| Northern Baltic | 1.00 | -0.29 | 10221.1 | 0.99 | -1.10 | 433.4 |
| Southern Baltic | 0.99 | -0.96 | 10151.9 | 1.00 | 0.34 | 439.7 |
| Carpathian | 0.99 | -1.24 | 10123.6 | 1.00 | -0.29 | 436.9 |
| Eastern European Lowlands | 0.98 | -2.03 | 10042.8 | 1.00 | 0.18 | 439.0 |
| Central European Lowlands | 0.98 | -1.74 | 10072.3 | 0.97 | -2.77 | 426.0 |
| Dinaric Balkan | 0.98 | -1.97 | 10049.2 | 1.00 | 0.13 | 438.8 |
| Northwestern Iberian | 0.98 | -2.23 | 10022.0 | 0.99 | -1.46 | 431.8 |
| Italian | 0.98 | -2.08 | 10037.3 | 0.96 | -4.17 | 419.9 |
| Average all European | | | 10212.2 | | | 436.4 |

**Table S7. Number of segregating (A) and fixed (B) deleterious mutations for moderate- and high-impact variants per population.** Ratio and percentage change relate each cluster to the average of all European wolves.

| A) Segregating | Moderate Impact Variants | | | High Impact Variants | | |
|---|---|---|---|---|---|---|
| Group | Ratio from average | Percent change from average | Segregating Deleterious | Ratio from average | Percent change from average | Segregating Deleterious |
| Italian | 0.55 | -44.61 | 15559 | 0.58 | -42.18 | 643 |
| Northwestern Iberian | 0.63 | -37.42 | 17578 | 0.64 | -36.42 | 707 |
| Scandinavian | 0.67 | -33.34 | 18725 | 0.68 | -31.65 | 760 |
| Western Finland | 0.89 | -11.04 | 24987 | 0.89 | -10.61 | 994 |
| Central European Lowlands | 0.95 | -5.24 | 26616 | 0.97 | -3.15 | 1077 |
| Carpathian | 1.01 | 0.80 | 28314 | 0.99 | -1.44 | 1096 |
| Eastern Finland | 1.05 | 5.00 | 29492 | 1.05 | 4.77 | 1165 |
| Dinaric Balkan | 1.17 | 17.47 | 32995 | 1.14 | 13.67 | 1264 |
| Southern Baltic | 1.16 | 15.83 | 32535 | 1.16 | 16.46 | 1295 |
| Northern Baltic | 1.20 | 20.21 | 33766 | 1.21 | 21.04 | 1346 |
| Karelian | 1.28 | 27.92 | 35930 | 1.28 | 27.79 | 1421 |
| Eastern European Lowlands | 1.44 | 44.41 | 40562 | 1.42 | 41.73 | 1576 |
| | | | 28088.3 | | | 1112 |

| B) Fixed | Moderate Impact Variants | | | High Impact Variants | | |
|---|---|---|---|---|---|---|
| Group | Ratio from average | Percent change from average | Fixed Deleterious | Ratio from average | Percent change from average | Fixed Deleterious |
| Italian | 3.54 | 253.60 | 293 | 2.75 | 175.41 | 14 |
| Northwestern Iberian | 2.81 | 181.19 | 233 | 2.75 | 175.41 | 14 |
| Scandinavian | 2.67 | 166.71 | 221 | 1.57 | 57.38 | 8 |
| Western Finland | 1.11 | 11.03 | 92 | 0.79 | -21.31 | 4 |
| Central European Lowlands | 0.74 | -26.38 | 61 | 0.59 | -40.98 | 3 |
| Carpathian | 0.72 | -27.59 | 60 | 0.79 | -21.31 | 4 |
| Eastern Finland | 0.58 | -42.07 | 48 | 0.59 | -40.98 | 3 |
| Dinaric Balkan | 0.42 | -57.76 | 35 | 0.59 | -40.98 | 3 |
| Southern Baltic | 0.49 | -50.52 | 41 | 0.39 | -60.66 | 2 |
| Northern Baltic | 0.41 | -58.97 | 34 | 0.39 | -60.66 | 2 |
| Karelian | 0.30 | -69.83 | 25 | 0.39 | -60.66 | 2 |
| Eastern European Lowlands | 0.04 | -96.38 | 3 | 0.39 | -60.66 | 2 |
| | | | 95.5 | | | 5.1 |

**References**

46. W. Zhang *et al.*, Hypoxia adaptations in the grey wolf (*Canis lupus chanco*) from Qinghai-Tibet Plateau. *PLOS Genet.* **10**, e1004466 (2014).
47. M.-H. S. Sinding *et al.*, Population genomics of grey wolves and wolf-like canids in North America. *PLOS Genet.* **14**, e1007745 (2018).
48. A. Auton *et al.*, Genetic recombination is targeted towards gene promoter regions in dogs. *PLOS Genet.* **9**, e1003984 (2013).
49. J. A. Robinson *et al.*, Genomic signatures of extensive inbreeding in Isle Royale wolves, a population on the threshold of extinction. *Sci. Adv.* **5**, eaau0757.
50. A. R. Perri *et al.*, Dire wolves were the last of an ancient New World canid lineage. *Nature* **591**, 87-91 (2021).
51. L. Loog *et al.*, Ancient DNA suggests modern wolves trace their origin to a Late Pleistocene expansion from Beringia. *Mol Ecol* **29**, 1596-1610 (2020).
52. M. Ní Leathlobhair *et al.*, The evolutionary history of dogs in the Americas. *Science* **361**, 81-85 (2018).
53. L. R. Botigué *et al.*, Ancient European dog genomes reveal continuity since the Early Neolithic. *Nat. Commun.* **8**, 16082 (2017).
54. S. Gopalakrishnan *et al.*, Interspecific gene flow shaped the evolution of the genus *Canis*. *Curr. Biol.* **28**, 3441-3449.e3445 (2018).
55. B. M. vonHoldt *et al.*, Whole-genome sequence analysis shows that two endemic species of North American wolf are admixtures of the coyote and gray wolf. *Sci. Adv.* **2**, e1501714 (2016).
56. Z. Fan *et al.*, Worldwide patterns of genomic variation and admixture in gray wolves. *Genome Res.* **26**, 163-173 (2016).
57. M. S. Sinding, S. Gopalakrishnan, K. Raundrup, L. Dalén, J. Threlfall, T. Gilbert, The genome sequence of the grey wolf, *Canis lupus linnaeus* 1758. *Wellcome Open Res.* **6**, 310 (2021).
58. L. A. Frantz *et al.*, Genomic and archaeological evidence suggest a dual origin of domestic dogs. *Science* **352**, 1228-1231 (2016).
59. A. H. Freedman *et al.*, Genome sequencing highlights the dynamic early history of dogs. *PLOS Genet.* **10**, e1004016 (2014).
60. M.-H. S. Sinding *et al.*, Arctic-adapted dogs emerged at the Pleistocene–Holocene transition. *Science* **368**, 1495-1499 (2020).
61. S. S. T. Mak *et al.*, Comparative performance of the BGISEQ-500 vs Illumina HiSeq2500 sequencing platforms for palaeogenomic sequencing. *GigaScience* **6**, 1-13 (2017).
62. L. Smeds, J. Aspi, J. Berglund, I. Kojola, K. Tirronen, H. Ellegren, Whole-genome analyses provide no evidence for dog introgression in Fennoscandian wolf populations. *Evol. Appl.* **14**, 721-734 (2021).
63. D. Yoo *et al.*, The genetic origin of short tail in endangered Korean dog, DongGyeongi. *Sci. Rep.* **7**, 10048 (2017).
64. D. Gómez-Sánchez *et al.*, On the path to extinction: Inbreeding and admixture in a declining grey wolf population. *Mol. Ecol.* **27**, 3599-3612 (2018).
65. J. Dabney *et al.*, Complete mitochondrial genome sequence of a Middle Pleistocene cave bear reconstructed from ultrashort DNA fragments. *Proc. Natl. Acad. Sci. U.S.A.* **110**, 15758-15763 (2013).

66. M. Meyer, M. Kircher, Illumina sequencing library preparation for highly multiplexed target capture and sequencing. *Cold Spring Harb. Protoc.* **2010**, pdb.prot5448 (2010).
67. C. Carøe *et al.*, Single-tube library preparation for degraded DNA. *Methods Ecol. Evol.* **9**, 410-419 (2018).
68. J. D. Kapp, R. E. Green, B. Shapiro, A fast and efficient single-stranded genomic library preparation method optimized for ancient DNA. *J. Hered.* **112**, 241-249 (2021).
69. M. P. Hoeppner *et al.*, An improved canine genome and a comprehensive catalogue of coding genes and non-coding transcripts. *PLOS ONE* **9**, e91172 (2014).
70. M. Schubert *et al.*, Characterization of ancient and modern genomes by SNP detection and phylogenomic and metagenomic analysis using PALEOMIX. *Nat. Protoc.* **9**, 1056-1082 (2014).
71. M. Schubert, S. Lindgreen, L. Orlando, AdapterRemoval v2: rapid adapter trimming, identification, and read merging. *BMC Res. Notes* **9**, 88 (2016).
72. H. Li, R. Durbin, Fast and accurate short read alignment with Burrows–Wheeler transform. *Bioinformatics* **25**, 1754-1760 (2009).
73. A. McKenna *et al.*, The Genome Analysis Toolkit: a MapReduce framework for analyzing next-generation DNA sequencing data. *Genome Res.* **20**, 1297-1303 (2010).
74. S. Rubinacci, D. M. Ribeiro, R. J. Hofmeister, O. Delaneau, Efficient phasing and imputation of low-coverage sequencing data using large reference panels. *Nat. Genet.* **53**, 120-126 (2021).
75. P. Danecek *et al.*, Twelve years of SAMtools and BCFtools. *GigaScience* **10**, giab008 (2021).
76. J. Marchini, B. Howie, Genotype imputation for genome-wide association studies. *Nat. Rev. Genet.* **11**, 499-511 (2010).
77. A. Manichaikul, J. C. Mychaleckyj, S. S. Rich, K. Daly, M. Sale, W.-M. Chen, Robust relationship inference in genome-wide association studies. *Bioinformatics* **26**, 2867-2873 (2010).
78. S. Purcell *et al.*, PLINK: a tool set for whole-genome association and population-based linkage analyses. *Am. J. Hum. Genet.* **81**, 559-575 (2007).
79. D. H. Alexander, J. Novembre, K. Lange, Fast model-based estimation of ancestry in unrelated individuals. *Genome Res.* **19**, 1655-1664 (2009).
80. V. Lefort, R. Desper, O. Gascuel, FastME 2.0: A comprehensive, accurate, and fast distance-based phylogeny inference program. *Mol. Biol. Evol.* **32**, 2798-2800 (2015).
81. N. Patterson, A. L. Price, D. Reich, Population structure and eigenanalysis. *PLOS Genet.* **2**, e190 (2006).
82. R. R. Fitak, OptM: estimating the optimal number of migration edges on population trees using Treemix. *Biol. Methods Protoc.* **6**, bpab017 (2021).
83. T. S. Korneliussen, A. Albrechtsen, R. Nielsen, ANGSD: analysis of next generation sequencing data. *BMC Bioinform.* **15**, 1-13 (2014).
84. A. Bergström *et al.*, Origins and genetic legacy of prehistoric dogs. *Science* **370**, 557-564 (2020).
85. J. Plassais *et al.*, Whole genome sequencing of canids reveals genomic regions under selection and variants influencing morphology. *Nat. Commun.* **10**, 1489 (2019).
86. P. Danecek *et al.*, The variant call format and VCFtools. *Bioinformatics* **27**, 2156-2158 (2011).

87. E. Garrison, Z. N. Kronenberg, E. T. Dawson, B. S. Pedersen, P. Prins, A spectrum of free software tools for processing the VCF variant call format: vcflib, bio-vcf, cyvcf2, hts-nim and slivar. *PLOS Comput. Biol.* **18**, e1009123 (2022).
88. C. L. Campbell, C. Bhérer, B. E. Morrow, A. R. Boyko, A. Auton, A pedigree-based map of recombination in the domestic dog genome. *G3 (Bethesda)* **6**, 3517-3524 (2016).
89. F. C. Ceballos, P. K. Joshi, D. W. Clark, M. Ramsay, J. F. Wilson, Runs of homozygosity: windows into population history and trait architecture. *Nat. Rev. Genet.* **19**, 220-234 (2018).
90. P. Cingolani *et al.*, A program for annotating and predicting the effects of single nucleotide polymorphisms, SnpEff: SNPs in the genome of *Drosophila melanogaster* strain w1118; iso-2; iso-3. *Fly (Austin)* **6**, 80-92 (2012).
91. G. Bertorelle *et al.*, Genetic load: genomic estimates and applications in non-model animals. *Nat. Rev. Genet.* **23**, 492-503 (2022).
92. D. Charlesworth, J. H. Willis, The genetics of inbreeding depression. *Nat. Rev. Genet.* **10**, 783-796 (2009).
93. R. Do, D. Balick, H. Li, I. Adzhubei, S. Sunyaev, D. Reich, No evidence that selection has been less effective at removing deleterious mutations in Europeans than in Africans. *Nat. Genet.* **47**, 126-131 (2015).
94. L. Laikre, N. Ryman, Inbreeding depression in a captive wolf (*Canis lupus*) population. *Conserv. Biol.* **5**, 33-40 (1991).
